# Bat-mediated pest regulation in understudied pasture agroecosystems

**DOI:** 10.64898/2026.06.04.730143

**Authors:** Hurpy Gwenaëlle, Aughney Tina, Moffat Aisling, Roche Niamh, Emma C. Teeling

**Affiliations:** School of Biology and Environmental Science, University College Dublin, Belfield, Dublin 4, Ireland; Bat Eco Services Limited, Virginia, Co. Cavan, Ireland; School of Agriculture and Food Science, University College Dublin, Belfield, Dublin 4, Ireland; Bat Conservation Ireland Carmichael House 4-7, North Brunswick Street, Dublin 7, Ireland

**Keywords:** agricultural pests, Chiroptera, Ireland, Tipula

## Abstract

Pastureland provides critical resources for livestock production. Despite the growing knowledge on pest regulating services provided by insectivorous bats, their role in pastureland systems remains understudied. We analysed the diets of four Irish bat species, *Plecotus auritus*, *Pipistrellus pygmaeus*, *Myotis nattereri*, and *Rhinolophus hipposideros*, to identify and categorise agricultural pest species based on metabarcoding studies in the pastureland rich agro-ecosystems of Ireland. Pest richness varied among bat species, with *P. auritus* consuming 22 known pest species, 13 in *P. pygmaeus*, 9 in *R. hipposideros*, and 7 in *M. nattereri*. Pest occurrence frequency varied in overall diets: *R. hipposideros* exhibited the highest (83%), followed by *P. auritus* (71%), *M. nattereri* (59%), and *P. pygmaeus* (32%). Importantly, 26 of the 35 pest species (74%) were consumed by only one bat species. Key pastureland pests, *Tipula oleracea* and *T. paludosa*, were detected across all bat species, with frequency of occurrence reaching up to 100% at several roosts. Livestock-associated disease vectors (*Culicoides scoticus* and *C. pulicaris*) were also consumed. Differences in diet among species reflect complementary foraging strategies, activity patterns, and habitat use, highlighting the collective role of bat communities in pest regulation. These findings provide additional evidence that insectivorous bats contribute to ecosystem services in pasturelands, potentially supporting reductions in pesticide use and enhancing sustainable pastureland management. By demonstrating the functional importance of bats beyond biodiversity conservation, this study underscores their value in nature-based solutions for sustainable agriculture and informs conservation and policy strategies in Ireland.

**Highlights:**

- Clear habitat bias in studies of ecosystem services by insectivorous bats.
- Bat species forage on diverse agricultural pests in pastureland-dominated areas.
- Bat foraging behaviour influences consumption of specific pest species.
- Bats consume major pasture pests, *Tipula oleracea* and *T. paludosa*.
- Different bat species target different pests, enhancing overall control.

## 1. Introduction

As global concerns about climate change and biodiversity loss intensify, improving the sustainability of agricultural systems has become a major priority. International initiatives such as the Convention on Biological Diversity (CBD, 1992), and the UN Decade on Ecosystem Restoration (2019), highlight the need to restore ecosystems and promote more sustainable land management. In Europe, the EU Biodiversity Strategy for 2030 (2020) and the Nature Restoration Law (2024) further emphasised the importance of enhancing biodiversity and restoring degraded ecosystems. In this context, strengthening natural pest regulation provided by insectivorous predators represents an important opportunity to support more sustainable agricultural practices.

The majority of the 1,500 bat species are insectivorous (Wilson and Mittermeier, 2019; Simmons and Cirranello, 2025). By foraging on a wide range of arthropods, insectivorous bats contribute to natural pest suppression in agricultural systems (Kunz et al., 2011; Russo et al., 2018; Ramírez-Fráncel et al., 2022). The diet of insectivorous bats includes numerous pest species such as leaf-feeding insects, livestock parasites, and vectors of human diseases (Ancillotto et al., 2017; Malso et al., 2017; Galan et al., 2018; Puig-Monserrat et al., 2020; Curran et al., 2022).

Through this predation, bats can reduce crop damage and support agricultural productivity and limit disease with their pest control services valued in the billions of dollars annually (Federico et al., 2008; Lopez-Hoffman et al., 2014; Maine et al., 2015; Wiederholt et al., 2015; Lopez-Hoffman et al., 2017; Rodriguez-San Pedro et., 2020).

Research into pest-regulating services provided by bats has produced substantial knowledge (Braun de Torrez et al., 2019; Linden et al., 2019; Rodriguez-San Pedro et al., 2020; Cohen et al., 2020; Montauban et al., 2021; Maslo et al., 2022; Hunninck et al., 2022; Bhalla et al, 2023b; Tuneu-Corral et al. 2023). However, this research has not been evenly distributed across ecological contexts, and certain ecosystems have remained overlooked. In addition, foraging differences among bat species and its effect on pest consumption may also be underrepresented in current research. Bat species exhibit a range of foraging behaviours (Jones and Rydell, 2005; Swartz et al., 2005; Denzinger and Schnitzler, 2013; McCracken et al., 2021; Russo, 2023). Such behavioural diversity may result in distinct ecosystem services, as variations in foraging behaviour influence the diversity and abundance of pest species consumed by bats (Ancillotto et al., 2017; Galan et al., 2018; Mata et al., 2021). Identifying gaps in current knowledge is essential for guiding future research efforts toward underrepresented habitats and bat species, rather than continuing to focus on well-studied ecosystems and bat–pest interactions. Addressing these gaps will improve our understanding of the variability of bat-mediated ecosystem services and strengthen their integration into conservation and management strategies (Fisher et al., 2009).

Ireland provides a particularly relevant context for investigating these potential ecosystem services. There are nine resident insectivorous bat species that occupy different foraging guilds and have been shown to play a role in pest suppression (Curran et al., 2022). Pastures cover approximately 61% of the country’s land area (Haughey et al., 2021), and agriculture plays a central role in the national economy. In 2023, dairy exports reached €6.5 billion, accounting for 34% of Ireland’s agri-food exports, while beef exports totaled €3.1 billion (Department of Agriculture, Food and the Marine, 2025). Cranefly larvae (*Tipula* spp.) are among the main pests affecting Irish pastures, reducing pasture yield through belowground herbivory that damages seedlings, roots, and young shoots (Blackshaw and Coll, 1999; Benefer et al., 2017; Moffat et al., 2022). Following the withdrawal of chlorpyrifos for agricultural use in 2019, effective chemical control options are limited, highlighting the importance of understanding natural pest regulation mechanisms. Importantly, *Tipula paludosa* and *T. oleracea* have been identified in bat diets (Galan et al., 2018; Mata et al., 2021; Curran et al., 2022). By preying on such pests, bats may contribute to maintaining pasture productivity and reducing reliance on chemical pesticides.

Here we firstly performed a scoping review to identify the gaps in knowledge of the ecosystem services provided by insectivorous bats from diverse foraging guilds across different habitats. Stimulated by these results, we identified pests consumed by four bat species with different foraging strategies (*Plecotus auritus*, *Pipistrellus pygmaeus*, *Rhinolophus hipposideros*, *Myotis nattereri*) in the overlooked pasture-dominated landscapes of Ireland, using published metabarcoding diet data (Hurpy et al., 2025). Finally we elucidated the bat-pest interactions and assessed the occurrence of two major pasture pests *- T. oleracea* and *T. paludosa*, highlighting the role of insectivorous bats in agricultural pest suppression.

## 2. Methods

### 2.1 Scoping review search

Following recommendations from Tricco et al. (2018), the scoping review was conducted by searching for publications on the ecosystem services provided by insectivorous bats. The review focused on (*a*) characterising the dominant habitat where the studies took place, (*b*) identifying the bat species studied, and (*c*) the pest species assessed. We carried out the search on the Web of Science (all databases) and Biological Science Index (Proquest) websites on the 10th of January 2024 using the following keywords: bat* AND pest suppression (Topic) or bat* AND pests suppression (Topic) or bat* AND ecosystem service* (Topic) not fruit (Topic) not frugivorous bat* (Topic).

To be included, the articles must have been peer reviewed and include the search keywords in the title, abstract or author’s keywords. A PRISMA systematic literature review flowchart guided the selection of studies included in this scoping review (Supplementary information Figure S1).

### 2.2 Data extraction from included studies

For each selected study, we extracted the year of sampling, geographic location (country and continent), and study scale (local, regional, or national). The dominant habitat was determined from the study description and classified following the habitat classification of the IUCN Red List (2024).

Details about the bat species studied were retrieved from included studies, as well as their foraging behaviour classified according to the classification proposed by Denzinger and Schnitzler (2013) (e.g. open space aerial forager, edge space aerial forager, edge space trawling forager, narrow space flutter forager, and narrow space passive gleaning forager). Any absent information was complemented by additional literature searches.

For studies reporting prey identification at the species level, lists of arthropod pest species consumed by bats were extracted. Pest species were classified according to their economic impact, including agricultural or forestry pests and species acting as vectors of human disease. For agricultural pests, the type of impact was further categorised (crop, plantation, pasture, or livestock). When pest status was not specified in the original studies, it was determined using the Arthemis database, the European and Mediterranean Plant Protection Organization (EPPO) Global Database, and, when relevant, additional local pest databases.

For studies reporting prey identification at the species level, lists of arthropod pest species consumed by bats were extracted. These records were used to identify bat-pest interactions, defined here as confirmed predation of an arthropod pest species by a bat species. To ensure reliable prey identification, only studies using faecal DNA metabarcoding were included in the bat–pest interaction network analysis.

### 2.3 Bat diet data

Metabarcoding data of bat faecal samples, collected from twelve maternity bat roosts in Ireland over a three-year period (2021–2023), were analysed from Hurpy et al. (2025) to identify the prey species remains present in the droppings. A total of 476,391,107 raw reads were obtained from NovaSeq sequencing, of which 255,022,827 remained after quality filtering. Taxonomic assignment resulted in 717 prey species, which were used for the current analyses. (Supplementary Information Tables S1 and S2).

The faecal samples were assigned to four bat species, *Plecotus auritus* (1,257 faeces), *Pipistrellus pygmaeus* (475), *Rhinolophus hipposideros* (168), and *Myotis nattereri* (44). These four bat species use distinct foraging strategies and exhibit different emergence patterns. *P. auritus* is primarily a narrow-space gleaning species, *M. nattereri* an edge-space forager combining aerial hawking and gleaning behaviour, *P. pygmaeus* an edge-space aerial hawker, and *R. hipposideros* a narrow-space aerial hawker. Species also differed in emergence timing, with *P. pygmaeus* and *R. hipposideros* emerging around sunset, whereas *P. auritus* and *M. nattereri* typically emerge later (Davidson-Watts and Jones, 2006; Denzinger and Schnitzler, 2013; Ancillotto and Russo, 2023; Jones and Froidevaux, 2023; Razgour et al., 2023; Schofield et al., 2023).

Land cover data surrounding each maternity roost were extracted from Hurpy et al. (2025). Improved grassland was the dominant land cover, ranging from 1.5% to 67% across sites. Six roosts were located in landscapes with more than 50% improved grassland, four roosts in landscapes with 5-50%, and two roosts in landscapes with less than 5%.

### 2.4 Arthropod prey pest status

To determine the pest status of arthropod prey, a list of pest species specific to Ireland was compiled by integrating data from both Irish and international sources. Irish sources consisted of official pest lists from the Department of Agriculture, Food and the Marine (2024), including the EU Priority Pests and Ireland’s Protected Zone Plant Pest lists. Additional information on pests affecting grasslands, crops, and livestock was obtained from Teagasc, the Agriculture and Food Development Authority (2017). Scientific literature was used to supplement sources (Colebrook and Wall, 2004; McCarthy et al., 2016; Piperaki and Daikos, 2016; Moffat et al., 2022; Nebbak et al., 2022). Pest risks to human health were addressed using data from the European Centre for Disease Prevention and Control (ECDC, 2024). The presence of each species in Ireland was verified using the National Biodiversity Data Centre database (National Biodiversity Data Centre, 2024). The resulting comprehensive pest list was cross-referenced with the list of prey species identified in the diets.

Climate change may alter pest distributions, enabling the establishment of new pest species in Ireland (Walther et al., 2009; Yan et al., 2017; Skendžić et al., 2021). Consequently, potential future pest species were also considered. We cross-referenced each prey species with two international agricultural pest databases, European and Mediterranean Plant Protection Organization (EPPO) Global Database (2024) and Arthemis database (INRAE-CBGP, 2024), to determine whether they are recognised as a pest in other regions. A final list of current and potential pest species was compiled, along with pest category assignments for each prey species identified at the species level in the bat diets.

For each pest species, one or more economic impact categories were assigned. Species were grouped into the following categories based on the sectors affected: Crop (arable and cereal crops), Pasture (grasslands used for grazing), Vegetables (horticultural and vegetable crops), Forestry (commercial forest plantations), Agroforestry (integrated tree–crop or tree–livestock systems), Livestock (direct impacts on farm animals), and Human Health (species acting as disease vectors or causing direct health impacts). Category assignments were based on information from official databases and the scientific literature.

### 2.5 Statistical analysis

Statistical analyses were conducted in R (version 4.3.3, R Core Team 2022) and RStudio (version 2023.12.1.402). For data visualisation, the *ggplot2* package (version 3.5.1 ; Wickham, 2016) was used.

Pest prey richness in each faecal sample was calculated using the ‘*hill_div*’ function from the *hilldiv* package (version 1.5.1), with the parameter q set to 0. The frequency of occurrence (FO) of each pest species in the diet was calculated for each bat species per sampling time point to determine how often pests were consumed. FO values were based on the total number of faecal samples analysed per bat species (Mata et al., 2021). Normality for both pest richness and FO was assessed using the ‘*shapiro.test’* function from the *stats* package (version 4.4.1). Due to the non-normal distribution of both richness and FO, a pairwise Wilcoxon-Mann-Whitney test was performed to compare pest richness per faecal sample and pest FO among bat species, year and reproductive period (Wilcoxon, 1947 ; Mann and Witney, 1947). For this, the ‘*compare_means’* function from *the ggpubr* package (version 0.6.3) was used (Kassambara, 2026).

Bat–pest interactions derived from the scoping review and from the Irish diet dataset were visualised using web plots created with the ‘*bipartiteD3*’ function from the *bipartiteD3* package (version 0.3.2; Terry, 2024). Difference in pest prey composition between bat species was tested using a permutational multivariate analysis of variance (PERMANOVA) using the ‘*adonis2*’ from vegan package with 999 permutations (Anderson et al., 2006; Anderson, 2014).

To characterise patterns of variation in the FO of *Tipula oleracea* and *T. paludosa* in the diets of the four bat species, comparisons between the FO of the two *Tipula* species were performed within each bat species using Wilcoxon–Mann–Whitney tests. FO values were calculated for each bat species overall and for each roost separately. *P. auritus*, *P. pygmaeus*, *M. nattereri*, and *R. hipposideros* were sampled at eight, four, three, and one roost respectively. Wilcoxon–Mann–Whitney tests were used to identify differences in FO between the two *Tipula* species for each bat species and at each roost. To examine relationships between the FO of *T. oleracea* and *T. paludosa* and surrounding grassland cover, the percentage of improved grassland around each bat roost was extracted from Hurpy et al. (2025), based on the Land Cover Map of Ireland. Since only *P. auritus* and *P. pygmaeus* were sampled across consecutive years and multiple roosts, landscape analyses were restricted to these species. Percentages of improved grassland cover were grouped into three categories: low (<5%), medium (5–50%), and high (>50%). Differences in FO among improved-grassland categories were tested using Kruskal-Wallis test (across all three categories) and Wilcoxon–Mann–Whitney pairwise comparison.

## 3 Results

### 3.1. Knowledge gaps identified by the scoping review

We identified 48 studies that met the inclusion criteria for the scoping review (Supplementary Information Table S3). The scoping review highlights a strong ecological bias in existing research on bat-mediated pest regulation, with a particular underrepresentation of pasture-dominated systems and limited representation of bat foraging guilds across studies. Habitats were strongly biased towards arable lands and plantations (34 studies), while pasture-dominated systems were comparatively understudied (only one study) (Figure1A). Across studies, foraging guild information was available for 66 species. Among these, most studies focused on edge-space aerial-hawking and open-space species (23 species each), with fewer studies on narrow-space flutter foragers (11 species) (Figure 1B). The list of bat species is provided in Supplementary Information Table S4.

**Figure 1.**
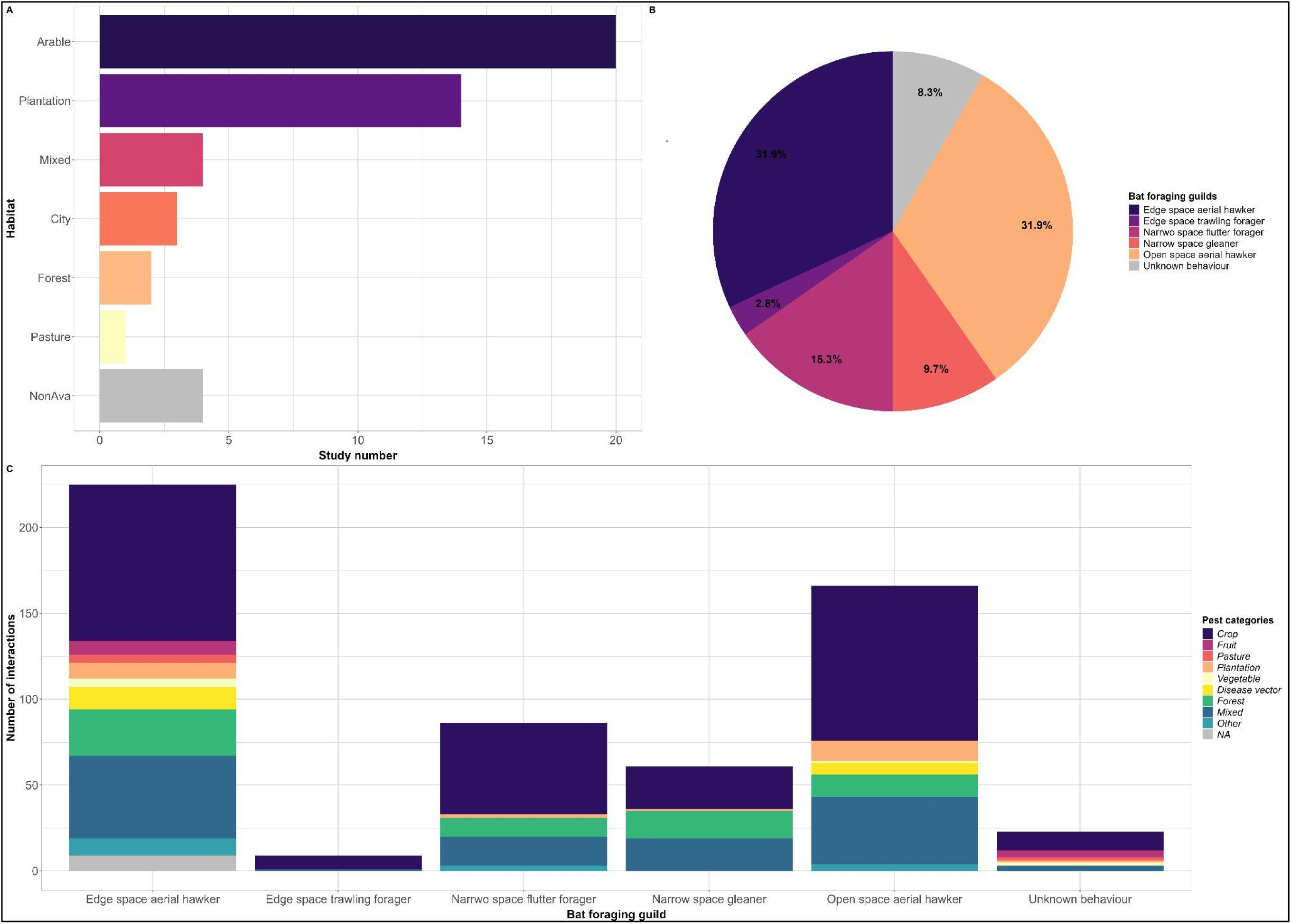
Representation of studies included in the scoping review. (A) Distribution of dominant habitat types across selected studies. (B) Proportion of bat foraging guilds represented across studies. (C) Number of recorded interactions between bat foraging guilds and pest categories. Agricultural pests include crop, fruit, pasture, plantation, and vegetable pests; Forestry pests include forest pests; Disease vectors include arthropod species acting as vectors of human diseases; Mixed refers to pests affecting multiple plant types; Other includes pest categories not assigned elsewhere; NA indicates unavailable information.

Analysis of pest records revealed interactions between bats and 149 arthropod pest species from Lepidoptera, Diptera, Hemiptera, and Coleoptera. Foraging guild patterns showed that edge-space and open-space aerial foragers interacted with the largest diversity of pest categories, preying on six and five categories, respectively. Agricultural pests were the most frequently consumed, particularly by open-space aerial (48% of interactions) and edge-space aerial foragers (46%). Forest pests were preyed upon by four guilds, dominated by edge-space aerial (37.5%) and narrow-space gleaning (33.3%) foragers. Interactions with insect pests that are vectors of human disease were restricted to edge-space and open-space aerial foragers (Figure 1C; Supporting Information Table S5).

Given the ecological importance of pasture landscapes in supporting both livestock production and bat foraging activity, these gaps limit our understanding of the functional role of bats in non-arable farming systems. To address this gap, we conducted a pest consumption-based analysis of insectivorous bats in pasture-dominated agricultural landscapes in Ireland, assessing their potential contribution to pest regulation.

### 3.2. Irish pest species in bat diets

Based on our novel list, 56 pest species are recognised as pests relevant to Ireland, although not all of these species are currently reported as present within the country (Supplementary Information Table S6).

In total, 35 of these arthropod pest species were detected in the diets of all four bat species; 71% of faecal samples from *Plecotus auritus*, 30% from *Pipistrellus pygmaeus*, 83% from *Rhinolophus hipposideros*, and 55% from *Myotis nattereri*.

Seven of these 35 pest species are considered as major pest species in Ireland and 28 are categorised as potential pests (Figure 2). These pest species belong to the orders Coleoptera, Diptera, and Lepidoptera, and were further grouped into six economic impact categories: Crop and Pasture, Crop and Vegetables, Forestry, Agroforestry, Livestock, and Human Health. Among these, 22, 13, 9, and 7 pest species were identified in the diet of *P. auritus*, *P. pygmaeus*, *R. hipposideros*, and *M. nattereri*, respectively (Figure 2). The number of pest categories detected per bat species ranged from three to six (Figure2).

**Figure 2.**
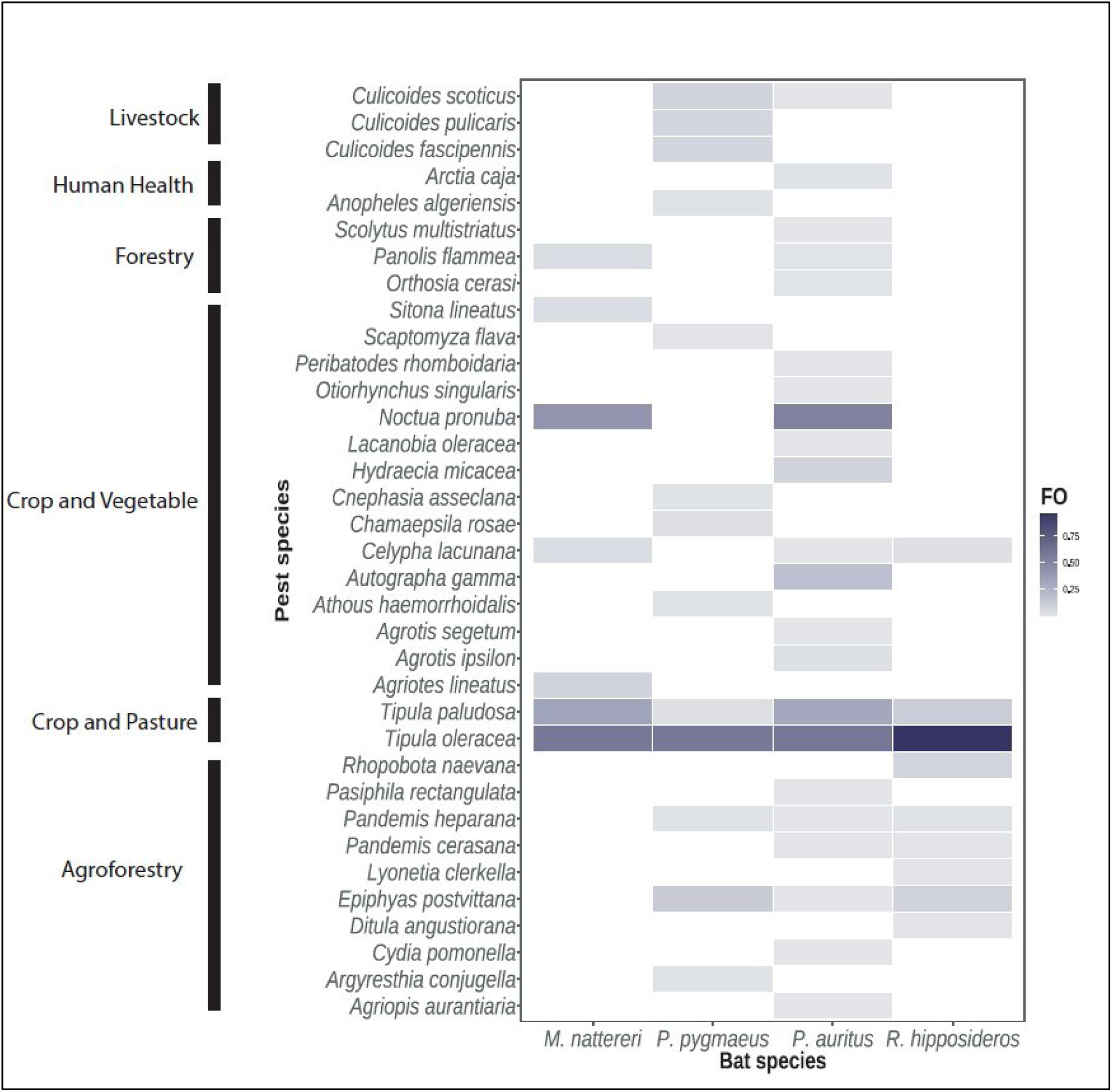
Frequency of occurrence (FO) of each pest species in the diet of each bat species. FO ranges from 0 to 1, representing the proportion of bat faecal samples in which each pest species was detected. Pest species are grouped by category.

The overall frequency of occurrence (FO) calculated across all pest species in each faecal sample per sampling event, ranged from 7.6% to 100%. *P. pygmaeus* exhibited the lowest mean FO (32.1%), while *P. auritus* and *R. hipposideros* showed the highest values (71.4% and 83.1%, respectively). *M. nattereri* showed intermediate FO values (mean 58.8%). Detailed FO values per year and sampling event are provided in Supplementary Figure S2. Comparison of mean FO between bat species revealed significant differences in FO between *P. pygmaeus* and both *P. auritus* and *R. hipposideros*, while a marginally significant difference was observed between *P. auritus* and *R. hipposideros* (Supplementary Table S7). FO of each pest species varied over the three-year study period. Several pest species exhibited high FO values (≥50%), including two major Diptera pest species, *Tipula paludosa* and *Tipula oleracea* (Craneflies), as well as potential pest Lepidoptera species, *Noctua pronuba* (Large yellow underwing) and *Rhopobota naevana* (Holly tortrix moth). Of the seven major pest species identified, five species were present in at least 5% of the faecal samples at one or more sampling periods. These included four Diptera species, *T. paludosa*, *T. oleracea*, *Culicoides pulicaris*, *C. scoticus* (biting midges), and one Lepidoptera species, *Epiphyas postvittana* (Light brown apple moth). The FO of individual pest species varied across the sampling period, with most species detected at one or a few sampling events. In contrast, three pest species (*Noctua pronuba*, *Tipula oleracea*, and *T. paludosa*) were detected across all sampling periods. (Supplementary Table S8).

Pest species richness per faecal sample showed variability among and within bat species across years and sampling periods. It ranged from zero to five pest species per sample, with mean values across bat species between 0.32 and 1.25. *P. auritus* exhibited highest mean richness (1.25 pest species per faecal sample) followed by *R. hipposideros* (1.08), *M. nattereri* (0.84), and *P. pygmaeus* (0.32) (Figure 3). Wilcoxon-Mann-Whitney tests indicated significant differences in pest richness between all four bat species (p-values ranging from 0.028 to < 2e-16), except between *P. auritus* and *R. hipposideros* (p = 0.250). Significant differences in pest richness across years were detected for *P. auritus* (p-value ranging from 0.016 to < 2e-16) and *P. pygmaeus* (p-value ranging from 0.0206 to 7.2e-6), while year-to-year differences were less pronounced for *M. nattereri* and *R. hipposideros*. When comparing pest richness across sampling periods, significant differences were detected between gestation and post-lactation (p-value = 7.3e-16), and between lactation and post-lactation (p-value = 0.002) for *P. auritus*, as well as between gestation and post-lactation for *R. hipposideros* (p-value = 0.002) (Supporting information Table S9).

**Figure 3.**
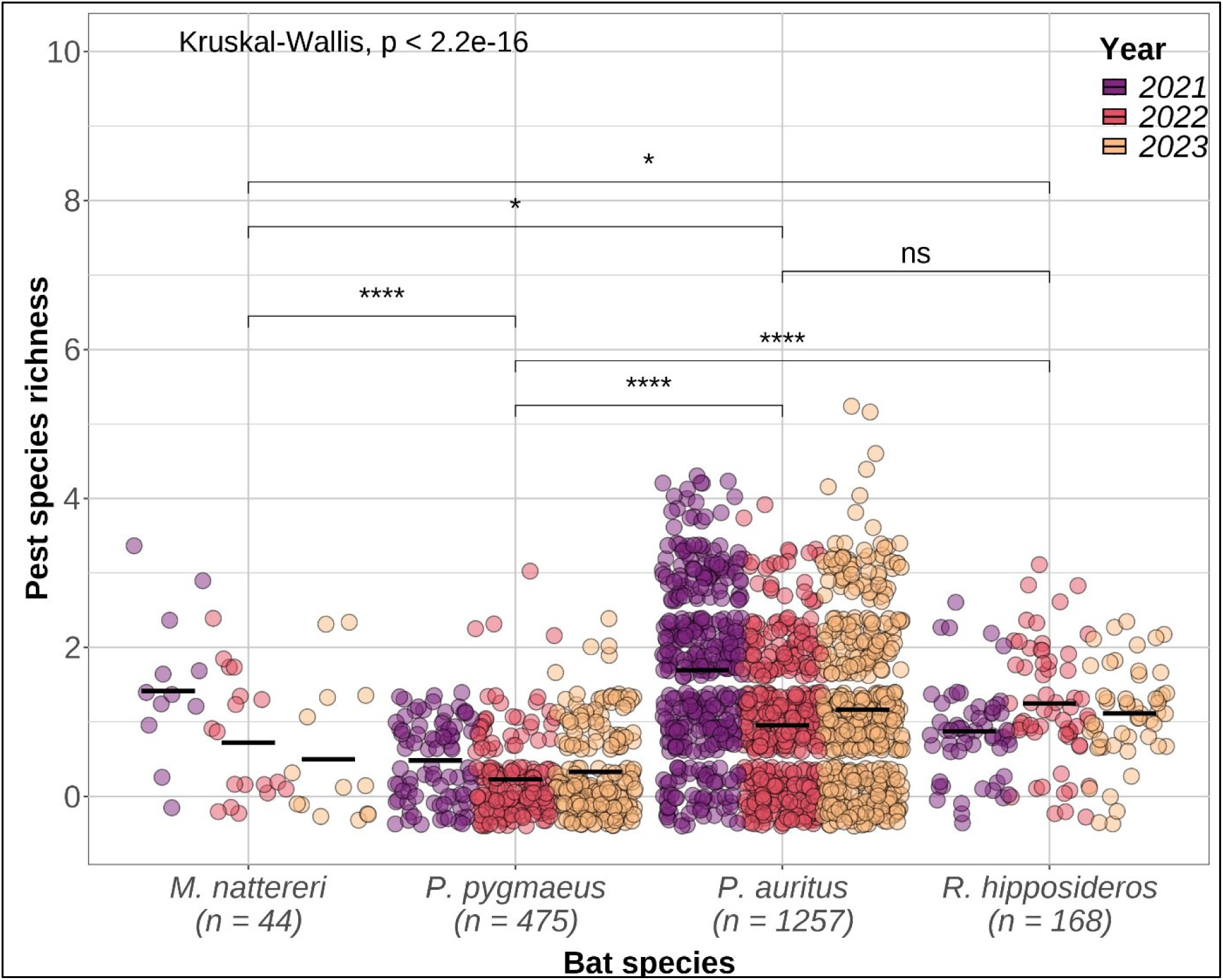
Jitter plot showing pest richness in faecal samples across years for each bat species. Each point represents a single faecal sample. Black bars indicate the median number of pest taxa per sample for each species and year. N is the total number of faecal samples analysed per bat species. Overall differences among bat species were assessed using the Kruskal-Wallis test (p < 2.2 × 10⁻¹⁶), as indicated on the plot. Pairwise comparisons between species were performed using the Wilcoxon-Mann-Whitney test; significance levels are indicated as follows: **** p < 0.0001, * p < 0.05, and ns, not significant

### 3.3. Bat pest species interactions

Results showed different patterns of species-specific interactions, with pest species identified in the diets of the four bat species, while other pests were consumed by individual bat species. Twenty-six pest species were found exclusively in the diets of specific bat species: 13 in *P. auritus*, eight in *P. pygmaeus*, three in *R. hipposideros,* and two in *M. nattereri* (Figure 2, Figure 4). Difference in pest prey composition between bat species was confirmed by the PERMANOVA analysis showing p-value < 0.001. (Supplementary Table S10).

**Figure 4.**
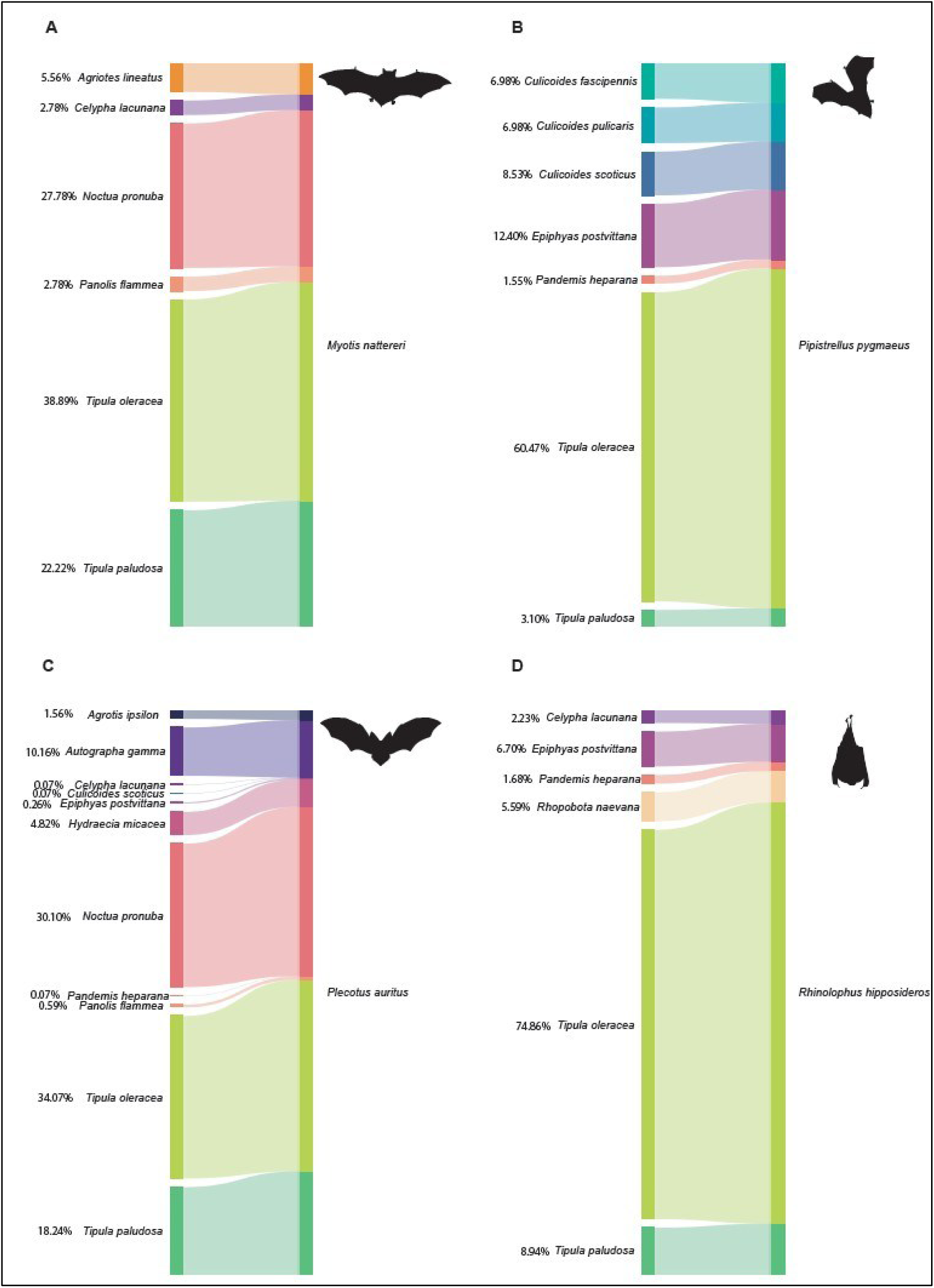
Interaction network between bat species and pest species. Only pest species with a frequency of occurrence exceeding 0.05 in at least one of the three years are shown. Panels: A, Myotis nattereri; B, Pipistrellus pygmaeus; C, Plecotus auritus; D, Rhinolophus hipposideros.

In terms of pest categories, all four bat species showed substantial interactions with crop and pasture pests, which accounted for between 51% and 82% of the total of interactions by bat species. Agroforestry pests were consumed primarily by *Rhinolophus hipposideros* (47%), with additional interactions from *Pipistrellus pygmaeus* (34%) and *P. auritus* (19%). Three pest categories - forestry, human health, and livestock pests - interacted with only two of the four bat species. Livestock pests were primarily consumed by *P. pygmaeus* (90%) and to a lesser extent by *P. auritus* (10%). Forestry pests were predominantly foraged by *P. auritus* (95%), with minor interactions from *Myotis nattereri* (5%). Similarly, pests categorised as human health threats (i.e. insect vectors of Human diseases) were mostly consumed by *P. auritus* (87.5%), with occasional interactions from the *P. pygmaeus* (12.5%).

Major pests, *T. oleracea* and *T. paludosa,* were both present in the diet of all four bat species, with FO values ranging from 1.2% to 100%, but with a higher FO in *P. auritus*, *N. nattereri*, and *R. hipposideros* than in the diet of *P. pygmaeus*. *C. pulicaris* presented FO values between 2% and 31%, whereas *C. scoticus,* which was found primarily in the diet of *P. pygmaeus*, and in the diet of *P. auritus* at one time collection point, showed FO values ranging from 0.7% to 6%. *E. postvittana* was found in the diet of *R. hipposideros*, *P. auritus* and *P. pygmaeus* with FO values ranging from 0.6% to 25% between gestation time points in 2022 and 2023 (Figure 4 and Figure 5).

**Figure 5.**
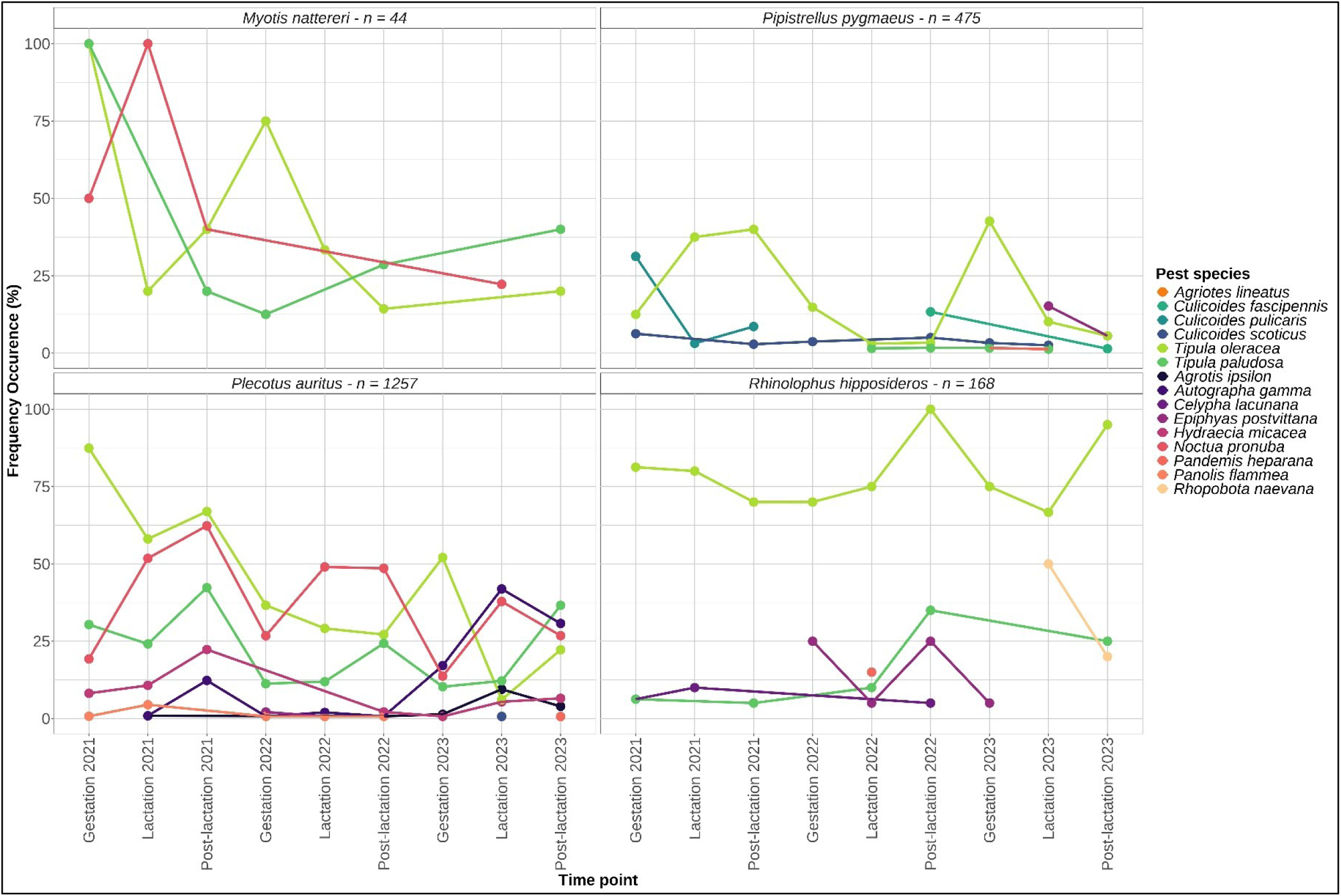
Frequency of occurrence (FO, %) of pest species in bat faecal samples across sampling events for each bat species. Only pest species with a frequency of occurrence exceeding 5% in at least one of the three years are shown.

*Plecotus auritus* exhibited the highest number of interactions with pest species, accounting for 82% of all recorded interactions, followed by *R. hipposideros* (9.6%), *P. pygmaeus* (6.9%), and *M. nattereri* (1.9%) (Supplementary Figure S2).

### 3.4. Major grassland pest, *Tipula oleracea* and *T. paludosa* in bat diets

Both *Tipula oleracea* and *T. paludosa* were detected in the diets of all four bat species. The combined frequency of occurrence (FO) of these species varied substantially among bats, with *Rhinolophus hipposideros* showing the highest mean FO (79.2%), followed by *Plecotus auritus* (48.5%), *Myotis nattereri* (44.3%), and *Pipistrellus pygmaeus* (18.6%). FO values were consistently high in *R. hipposideros* (minimum FO 66.7%), whereas values were more variable in *M. nattereri*, *P. auritus*, and *P. pygmaeus*, ranging from 0% to 100% (Supplementary Table S11).

Across all bat species, *T. oleracea* generally occurred more frequently than *T. paludosa*. Significant differences in FO between the two species were observed in *P. pygmaeus* (p = 2.8E10⁻⁴), *P. auritus* (p = 5.7E10⁻⁵), and *R. hipposideros* (p = 3.8E10⁻⁴), whereas no significant difference was detected for *M. nattereri* (p ≥ 0.35) (Figure 6). *P. pygmaeus* showed the lowest FO overall, with median values of 5% for *T. oleracea* and 0% for *T. paludosa*. FO values also varied among roosts, particularly for *P. auritus*, where some roosts showed marked differences between the two Tipula species while others did not (Figure 7). In contrast, FO values in *R. hipposideros*, sampled at a single roost (Kinvarra), showed *no overlap* between *T. oleracea* and *T. paludosa*, indicating a consistent pattern across sampling periods. Within species, FO values also varied across roost sites and sampling periods. For each bat species at various sampling periods, *Tipula* pest species were identified in all faecal samples, with FO values reaching up to 100%. For *R. hipposideros*, which was sampled exclusively from a single roost, *Tipula* pest species consistently occurred across all sampling periods, with FO values ranging from 66.67% to 100%. For *P. pygmaeus*, samples from the Birr roost showed high FO values for *Tipula* species, ranging from 7.69% to 100%, with an average FO of 52.09%. Similarly, *P. auritus* exhibited maximum FO values of 100% for *Tipula* species across six roosts. *M. nattereri* also demonstrated high FO values, with a maximum of 100% observed at the Glengariff roost. (Figure 7, Supplementary Information Table S11).

**Figure 6.**
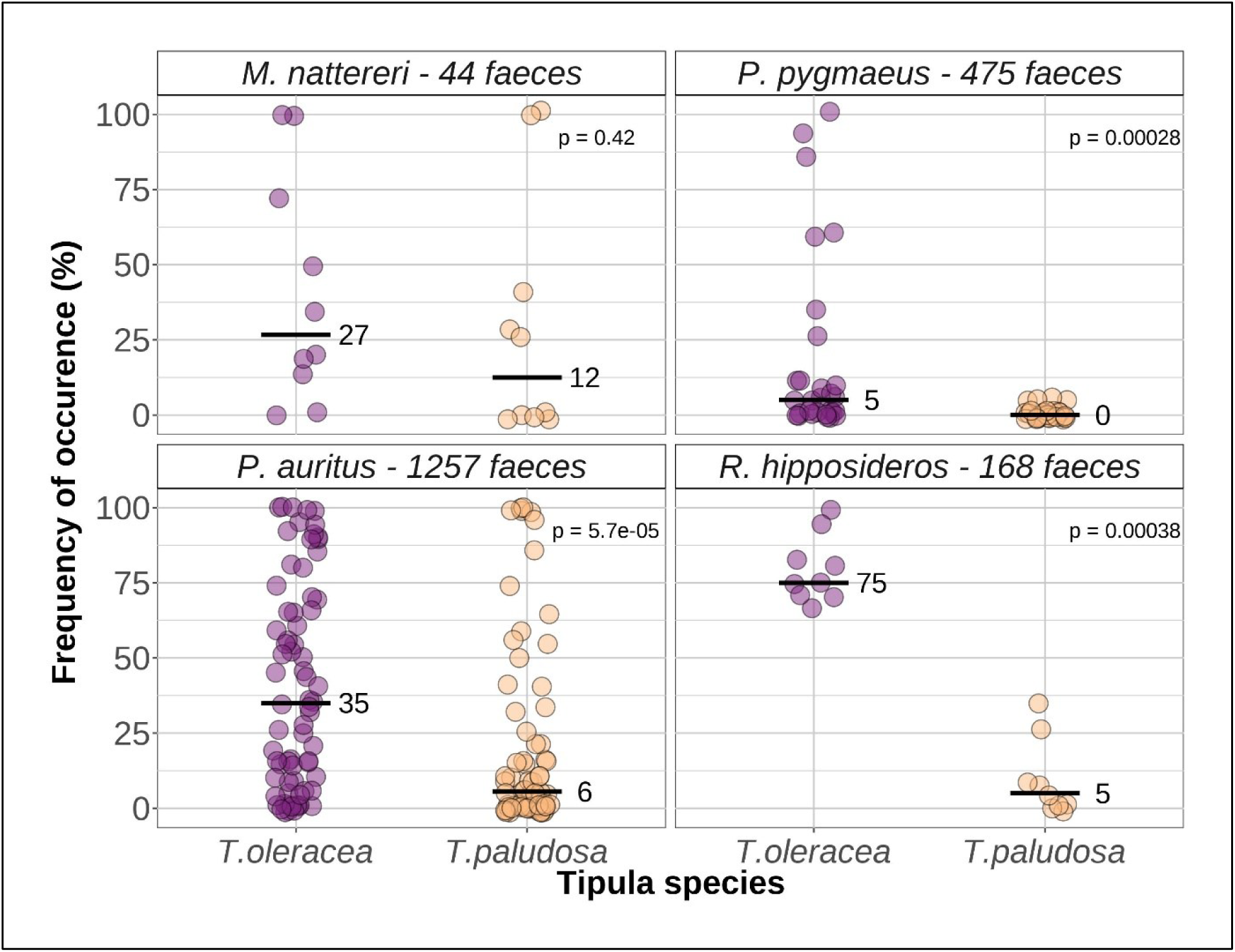
Frequency of occurrence (FO, %) of Tipula oleracea (purple dots) and T. paludosa (orange dots) at each sampling event, shown for each bat species. Crossbars indicate median values (reported on the right). P values from Wilcoxon–Mann–Whitney tests comparing FO between the two Tipula species are shown for each bat species.

**Figure 7.**
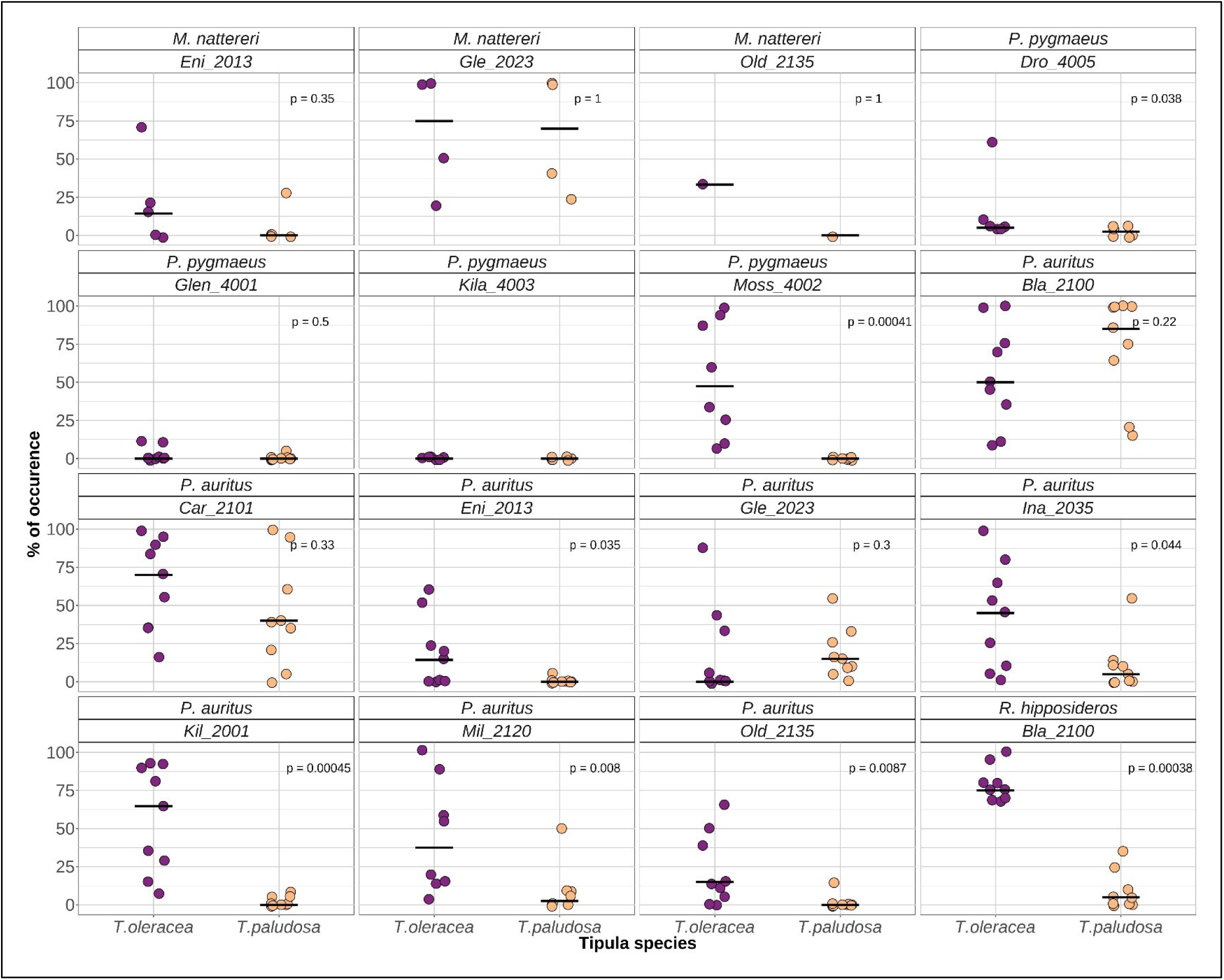
Frequency of occurrence (FO, %) of Tipula oleracea (purple dots) and T. paludosa (orange dots) at each sampling event, shown for each roost and bat species. Crossbars indicate median values. P values from Wilcoxon–Mann–Whitney tests comparing the mean FO between the two Tipula species are shown for each roost

To assess whether landscape context influenced the occurrence of major grassland pests in bat diets, the relationship between improved grassland cover surrounding roosts and the FO of *Tipula* species was examined for *P. auritus* and *P. pygmaeus*. For *P. auritus*, the frequency of occurrence (FO) of *Tipula oleracea* differed significantly depending on improved grassland cover percentages surrounding roosts (Kruskal–Wallis test, p = 0.0082; Figure 8). FO was significantly higher at roosts where improved grassland cover exceeded 50% compared to roosts with less than 5% improved grassland cover (p-value = 0.0022). No significant difference in FO was detected between low (<5%) and medium (5–50%) improved grassland cover classes (Figure 8). For *Pipistrellus pygmaeus*, FO of *T. oleracea* was significantly higher at roosts surrounded by high (>50%) compared to medium (5–50%) improved grassland cover (p = 0.00015; Figure 8). In contrast, FO of *Tipula paludosa* did not vary significantly with the proportion of improved grassland cover for both *P. auritus* and *P. pygmaeus*.

**Figure 8.**
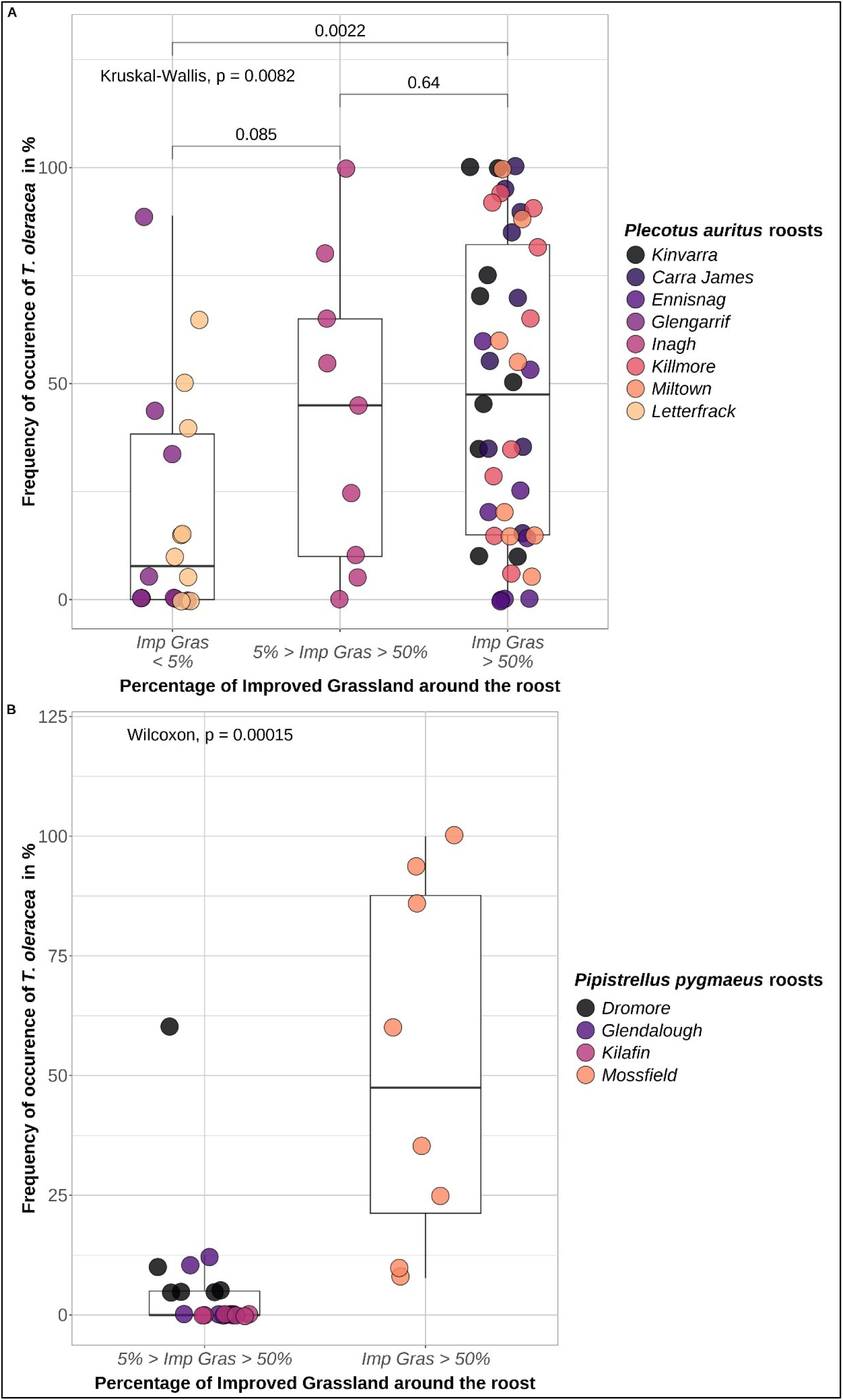
Frequency of occurrence (FO, %) of Tipula oleracea at each bat roost and sampling event across categories of improved grassland cover (Imp. grassland: low <5%, medium 5–50%, high >50%) for Plecotus auritus (A) and Pipistrellus pygmaeus (B). P values from Kruskal–Wallis and Wilcoxon–Mann–Whitney pairwise comparisons are shown for P. auritus, and from Wilcoxon–Mann–Whitney tests for P. pygmaeus.

## 4. Discussion

The scoping review highlighted critical gaps in the current knowledge of pest-regulation services provided by insectivorous bats within agricultural systems, particularly in pasture-dominated landscapes. This gap is notable considering that pasturelands cover approximately 25% of the Earth’s surface (FAO, 2025), provide the primary forage base for livestock production and play an essential role in supporting dairy and meat production (Bengtsson et al., 2019; Fraser et al., 2022). Rectifying this knowledge gap is important given that pasture pests are known to reduce productivity, with cascading effects on livestock production (Jackson et al., 2012; Moffat et al., 2022). In addition, blood-feeding insects negatively affect animal welfare, reduce productivity, and can transmit fatal diseases (Wilson and Mellor, 2009; Kazek and Jezierski, 2014).

In this study, we address this gap by investigating pest consumption by insectivorous bats in pasture-dominated agricultural landscapes in Ireland.

### 4.1. Pest Richness in bat diets

This study demonstrates that Irish bat species exploit a diverse assemblage of agricultural pests across multiple taxonomic groups, highlighting their underappreciated role in pastureland ecosystems. Pest consumption patterns observed in *P. auritus, P. pygmaeus,* and *M. nattereri* are consistent with those reported in other European studies, confirming broadly similar dietary patterns across habitats (Shiel et al., 1991; Mata et al., 2021; Puig-Monserrat et al., 2020; Ancillotto et al., 2022; Razgour et al., 2023). In *R. hipposideros*, the detection of key agricultural pests such as *Epiphyas postvittana* and *Tipula paludosa* represents, to our knowledge, the first record of these species in its diet in Ireland, providing new evidence of its contribution to pest suppression.

Despite lower pest richness than in some studies (Mata et al., 2021; Curran et al., 2022; Miñarro et al., 2026), frequency of occurrence (FO) values were high across all bat species, ranging from 32% in *P. pygmaeus* to 83% in *R. hipposideros*. This indicates that bats consistently exploit pest species when available, even when overall pest diversity is limited. This also suggests that frequency of occurrence may be a more relevant indicator of pest suppression potential than richness alone, as it reflects the consistency of bat-pest interactions rather than simply dietary breadth.

Variation in pest richness across studies is widely reported in the literature and likely reflects multiple ecological and methodological factors. As highlighted by Mata et al. (2021), sampling across diverse habitats can increase the number of pest species detected. In the present study, many sampling sites were dominated by improved grassland, with relatively limited habitat diversity, which may have constrained the range of available pest species. In addition, Ireland supports lower insect diversity compared to other European regions (McCarthy, 1986; Ferriss, 2009; Harrison, 2014), which may further contribute to reduced pest richness. Methodological factors may also play a role, as conservative metabarcoding thresholds can exclude rare prey items and lead to underestimation of dietary diversity (Alberdi et al., 2018), as well as the limited number of faecal samples from *M. nattereri*.

Finally, variation in pest richness and occurrence across bat species, sampling periods, and locations likely reflects temporal and spatial differences in prey availability, including seasonal emergence patterns and interannual fluctuations in insect populations (Davies et al., 2022; Moffat et al., 2022). However, these patterns are also shaped by species-specific differences in prey use and foraging behaviour, highlighting the role of bat and prey traits in structuring bat–pest interactions across agricultural landscapes (Puig-Monserrat et al., 2020; Miñarro et al., 2026).

### 4.2. Bat and Prey Traits Drive Differences in Pest Exploitation among Species

The four bat species investigated in this study showed clear differences in the pest species detected in their diets, with limited overlap (26%), indicating complementary exploitation of the pest community. These patterns likely reflect contrasts in prey accessibility shaped by foraging strategy, emergence timing, habitat, and prey traits, which together influenced prey selection (Nguyen et al. 2019; Puig-Montserrat et al., 2020).

Foraging strategy, activity timing, and pest behaviour strongly influence which pests are accessible to each species. Differences in the consumption of *Tipula oleracea* and *T. paludosa* illustrate this pattern. While both species emerge from the soil, *T. paludosa* females tend to remain on the grass with limited flight ability, whereas *T. oleracea* females are capable of sustained flight over longer distances and remain active throughout the night (Robertsoon, 1939; Blackshaw and Coll, 1999; Blackshaw and Petrovskii, 2007; Petersen, 2013). Consequently, *T. paludosa* is more accessible to gleaning or clutter-adapted foragers (*P. auritus* and *M. nattereri*), whereas *T. oleracea* is more available to aerial-hawking species (*R. hipposideros* and *P. pygmaeus*).

A similar pattern is observed for temporal alignment between bat activity and prey behaviour. In our study, predation on *Culicoides* species (e.g., *C. scoticus* and *C. pulicaris*) was primarily associated with *P. pygmaeus*. These two species and the majority of Culicoides species are crepuscular species and swarm in open space close to breeding sites (Viennet et al., 2012 ; Meiswinkel and Elbers, 2016 ; Mullen and Murphree, 2019). This behaviour aligns well with the early emergence time and foraging behaviour of *P. pygmaeus*. In contrast, later-emerging and clutter-adapted bat species (Ancillotto and Russo, 2023; Jones and Froidevaux, 2023; Razgour et al., 2023; Schofield et al., 2023), show little evidence of consuming these species, likely reflecting differences in activity timing and foraging niches.

By exploiting pests that vary in space, time, and behaviour, bat species collectively target a broader assemblage than any single species alone, highlighting their complementary roles in pest suppression. This complementarity suggests that bat communities may enhance the consistency and overall effectiveness of pest regulation in pastureland systems.

### 4.3. Importance of ecosystem services provided for pastureland

The presence of two major pastureland pests, *T. oleracea* and *T. paludosa*, in the diets of all four bat species demonstrates that bats are consistently exploiting major pests of pastureland productivity. Larvae of these two Tipulidae species reduce pasture yield through belowground herbivory, damaging seedlings, root systems and young shoots (Blackshaw and Coll 1999; Benefer et al., 2017), and directly affect pastureland yield and by consequence livestock feed availability (Moffat et al., 2022). The importance of this ecosystem service is further emphasised by the ban on chlorpyrifos, previously used to control these pests (Commission of Non-renewal of the approval of the active substance chlorpyrifos, 2020).

The consistent detection of *Tipula* spp. across bat species and sampling periods, together with the positive relationship between their occurrence in bat diets and the proportion of surrounding grassland, suggests that bats regularly exploit these pests in response to local availability.

Bats are not the only insectivorous vertebrates contributing to the regulation of Tipulidae populations. Ground-feeding birds, such as Dunlin (*Calidris alpina pacifica*, European golden plover (*Pluvialis apricaria*), Oystercatchers (*Haematopus ostralegus*), Red-billed Chough (*Pyrrhocorax pyrrhocorax*) (McCracken et al., 1992; Zwarts and Blomert, 1996; Evans-Ogden et al., 2008; Pearce-Higgins et al., 2010) have also been reported to consume crane flies, during larval or adult emergence periods. However, bat predation complements these diurnal predators by targeting adult pests at night, providing continuous, round-the-clock predation pressure. This temporal complementarity may strengthen the overall effectiveness of biological pest control in pastureland landscapes.

Beyond Tipulidae, *P. pygmaeus* consumed *Culicoides* midges (*C. scoticus* and *C. pulicaris*), which are vectors of livestock diseases such as Schmallenberg and Bluetongue viruses (Mullen and Murphree, 2019). This suggests an additional ecosystem service through the potential regulation of disease vectors. In the context of climate change, which is expected to alter midge distributions and increase the risk of vector-borne diseases in Ireland (Wittmann and Baylis, 2000; Walther et al., 2009; Brand and Keeling, 2017; Yan et al., 2017; Skendžić et al., 2021; Hurpy et al., 2025), this role may become increasingly important.

Taken together, these findings show that bat communities contribute to the suppression of key pastureland pests and livestock disease vectors, supporting more productive and resilient pastureland systems.

### 4.4. Conservation Implication, Policy Relevance and Future Research

Bats contribute to pest regulation in pastureland systems, reinforcing their functional importance beyond biodiversity conservation. By consuming key agricultural pests, bat communities represent a component of nature-based solutions supporting sustainable grassland management. These findings are particularly relevant in a policy context marked by reduced chemical pest control and increasing recognition of ecosystem service-based solutions (EU Biodiversity Strategy for 2030, 2020; Nature Restoration Law, 2024; Sustainable Development Goal 15, 2025). As agricultural systems shift toward lower pesticide reliance, maintaining wildlife-mediated pest regulation becomes increasingly important.

In Ireland, pasture-based agriculture dominates land use and is a cornerstone of the national economy (Annual Review and Outlook for Agriculture, Food, and the Marine 2024–2025, 2025). Given its economic significance, the government has set ambitious sustainability targets through the Food Vision 2030 strategy (Food Vision 2030, 2021), aiming to create a “Climate Smart, Environmentally Sustainable Agri-Food Sector” by increasing organic farming to 7.5% of agricultural land, prioritising biodiversity on 10% of farmed land, and reducing pesticide use by 50% by 2030. In this pasture-dominated system, pests affecting grassland productivity and livestock health have direct implications for food security and the economy. Bats contribute ecosystem services that align directly with these national objectives, offering nature-based pest regulation that supports sustainable pasture management.

Quantifying the economic value of bat-mediated pest suppression remains a critical step for understanding their role in pasture-based systems. Future research should link bat diet data with Tipula population dynamics to determine whether predation reduces pest abundance during key emergence periods, and assess the consequences of larval predation on pastureland productivity. In this study, the sampling focused on bat reproductive periods, aligning future research with Tipula emergence would improve our understanding of prey availability and bat-pest interactions. The two species exhibit distinct life histories, with *T. paludosa* being univoltine and emerging in late summer, whereas *T. oleracea* is bivoltine with emergence peaks in spring and autumn (Moffat et al., 2022). Additionally, expanding such studies to additional bat species and pastureland systems will clarify the collective contribution of bat communities to pastureland productivity and provide evidence to support sustainable agricultural policies in Ireland.

Maintaining these ecosystem services depends on effective conservation of bat populations. This includes protecting maternity roosts, preserving commuting routes and foraging habitats, and reducing pressures such as artificial light at night (Russo et al., 2024). Integrating bat conservation into agri-environment schemes, including support for roost creation and habitat management, could strengthen the contribution of bats to sustainable pastureland systems while supporting broader biodiversity objectives. Policy measures, including agri-environment schemes that fund roost creation and habitat enhancement, could further strengthen bat conservation while simultaneously supporting pest suppression.

## 5. Conclusion

Pest-regulating services provided by bat species in pastureland context remain a key knowledge gap in agricultural ecosystems. This study provides new evidence of the role of four Irish bat species in consuming key pastureland pests and livestock disease vectors. These results add to growing evidence that insectivorous bats deliver substantial ecological and economic benefits by reducing pest abundance and limiting reliance on pesticides. Our findings reinforce the functional importance of bats beyond biodiversity conservation and highlight their contribution to sustainable pastureland management. Furthermore, the study strengthens the case for integrating bat conservation into agricultural policy frameworks and for recognising bats as vital providers of ecosystem services in Ireland.

## Acknowledgement

This research was supported by the Irish Research Council of Ireland, Postgraduate Scholarship (grant no. GOIPG/2020/1517) awarded to G.H. and E.C.T. and National Parks and Wildlife Service (NPWS) of Ireland funding awarded to E.C.T.

## Data Availability Statement

Data supporting this article will be available after publication of the manuscript.

