## Supplementary Information for "Bat-mediated pest regulation in understudied pasture agroecosystems"

**Affiliation addresses**

[1]School of Biology and Environmental Science, University College Dublin, Belfield, Dublin 4, Ireland.

[2]Bat Eco Services Limited, Virginia, Co. Cavan, Ireland.

[3]School of Agriculture and Food Science, University College Dublin, Belfield, Dublin 4, Ireland.

[4]Bat Conservation Ireland Carmichael House 4-7, North Brunswick Street, Dublin 7, Ireland

**Corresponding author**

*Emma Teeling:

**Table of contents**

| | |
|---|---|
| Page 2 | Figure S1 |
| Page 3 | Table S3 |
| Page 9 | Table S4 |
| Page 16 | Table S6 |
| Page 21 | Figure S2 |
| Page 22 | Table S7 |
| Page 23 | Table S8 |
| Page 27 | Table S9 |
| Page 28 | Table S10 |
| Page 29 | Table S11 |

***Figure S1.*** *PRISMA flowchart used to select studies included in the scoping review for insectivorous bats and ecosystem services assessment*

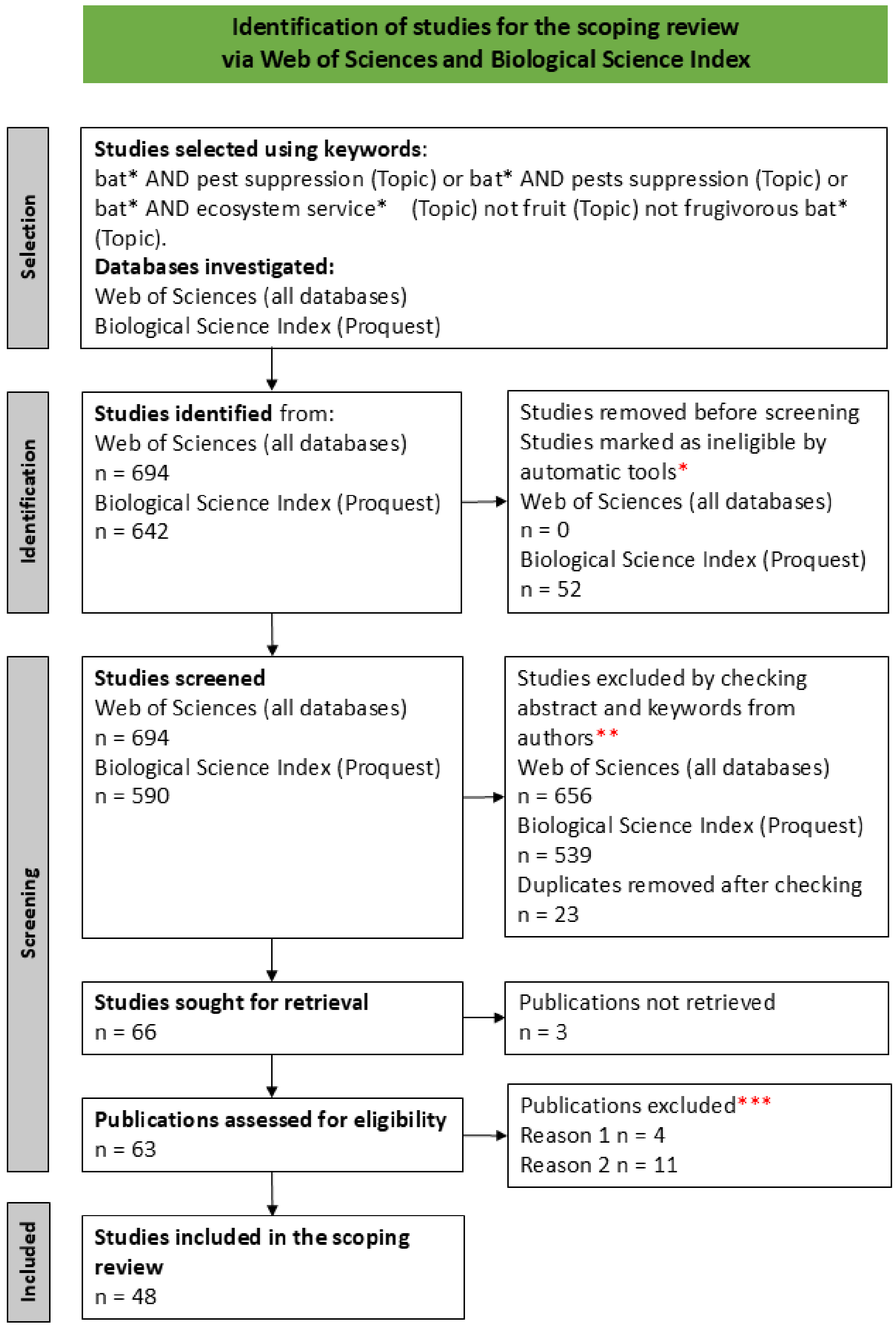

* Publications included only articles published in scholarly journals and written in English.

** Excluded publications that do not assess the ecosystem services provided by insectivorous bats.

*** Reason 1: ecosystem services assessment from bats were not separately evaluated of other insectivorous predators; Reason 2 : ecosystem services not evaluated.

***Table S3***. *List of selected studies for the scoping review. For each study information is provided: country where the study took place, main habitat, and bat species investigated. For visibility, if the study investigated more than three bat species, "Bat assemblage" is indicated.*

| Study nb | Authors | Article title | Country | Main habitat type | Bat species |
|---|---|---|---|---|---|
| 1 | Aguiar L et al. (2021) | Going out for dinner-The consumption of agriculture pests by bats in urban areas | Brazil | Urban area | Bat assemblage |
| 2 | Aihartza J et al. (2023) | Aerospace-foraging bats eat seasonably across varying habitats | Spain | Arable land | *Miniopterus schreibersii* |
| 3 | Aizpurua O et al. (2018) | Agriculture shapes the trophic niche of a bat preying on multiple pest arthropods across Europe: Evidence from DNA metabarcoding | Europe | No information | *Miniopterus schreibersii* |
| 4 | Ancillotto L et al. (2017) | Effects of free-ranging cattle and landscape complexity on bat foraging: Implications for bat conservation and livestock management | Italy | Pastureland | Bat assemblage |
| 5 | Ancillotto L et al. (2022) | Bats as suppressors of agroforestry pests in beech forests | Italy | Forest | *Barbastella barbastellus*<br>*Plecotus auritus* |
| 6 | Baroja U et al. (2021) | Bats actively track and prey on grape pest populations | Spain | Arable land | Bat assemblage |
| 7 | Beilke and O'Keefe (2023) | Bats reduce insect density and defoliation in temperate forests: An exclusion experiment | US | Forest | Bat assemblage |
| 8 | Bhalla et al. (2023a) | Batting for rice: The effect of bat exclusion on rice in North-East India | India | Arable land | Bat assemblage |
| 9 | Bhalla et al (2023b) | Insectivorous bats in Indian rice fields respond to moonlight, temperature, and insect activity | India | Arable land | Bat assemblage |
| 10 | Bouarakia et al. (2023) | Bats and birds control tortricid pest moths in South African macadamia orchards | South Africa | Plantation | Bat assemblage |

| Study nb | Authors | Article title | Country | Main habitat type | Bat species |
|---|---|---|---|---|---|
| 11 | Braun de Torrez et al. (2019) | Sympatric Bat Species Prey Opportunistically on a Major Moth Pest of Pecans | Texas | Plantation | Bat assemblage |
| 12 | Burgar et al. (2021) | Effectiveness of bat boxes for bat conservation and insect suppression in a Western Australian urban riverine reserve | Australia | Urban area | *Chalinolobus gouldii* |
| 13 | Charbonnier et al. (2021) | Pest control services provided by bats in vineyard landscapes | France | Arable land | Bat assemblage |
| 14 | Cohen et al. (2020) | An appetite for pests: Synanthropic insectivorous bats exploit cotton pest irruptions and consume various deleterious arthropods | Israel | Plantation | *Pipistrellus kuhlii* |
| 15 | Curran et al. (2022) | One bat's waste is another man's treasure: a DNA metabarcoding approach for the assessment of biodiversity and ecosystem services in Ireland using bat faeces | Ireland | No information | *Rhinolophus hipposideros* |
| 16 | Da Silva et al. (2023) | Can bats help paper industry? An evaluation of eucalypt insect-related predation by bats | Portugal | Plantation | Bat assemblage |
| 17 | Davidai et al. (2015) | The importance of natural habitats to Brazilian free-tailed bats in intensive agricultural landscapes in the Winter Garden region of Texas, United States | USA | Arable land | *Tadarida brasiliensis* |
| 18 | Federico et al. (2008) | Brazilian free-tailed bats as insect pest regulators in transgenic and conventional cotton crops. | USA | Plantation | *Tadarida brasiliensis* |
| 19 | Harms et al. (2020) | Investigating Bat Activity in Various Agricultural Landscapes in Northeastern United States | USA | Arable land | Bat assemblage |

| Study nb | Authors | Article title | Country | Main habitat type | Bat species |
|---|---|---|---|---|---|
| 20 | Hughes et al. (2021) | Big bats binge bad bugs: Variation in crop pest consumption by common bat species | USA | Mixed | Bat assemblage |
| 21 | Hunninck et al. (2022) | Far from home: Bat activity and diversity in row crop agriculture decreases with distance to potential roost habitat | USA | Arable land | Bat assemblage |
| 22 | Karp and Daily (2014) | Cascading effects of insectivorous birds and bats in tropical coffee plantations. | Costa Rica | Plantation | Bat assemblage |
| 23 | Karp et al. (2013) | Forest bolsters bird abundance, pest control and coffee yield | Costa Rica | Plantation | Bat assemblage |
| 24 | Kemp, et al. (2019) | Bats as potential suppressors of multiple agricultural pests: A case study from Madagascar | Madagascar | Mixed | Bat assemblage |
| 25 | Linden et al. (2019) | Ecosystem services and disservices by birds, bats and monkeys change with macadamia landscape heterogeneity | South Africa | Plantation | Bat assemblage |
| 26 | Liu et al. (2024) | Pest suppression services and dietary niche differentiation of bats in Chinese smallholder farming systems: implications for integrated pest management | China | Urban area | Bat assemblage |
| 27 | Lopez-Hoffman et al. (2014) | Market Forces and Technological Substitutes Cause Fluctuations in the Value of Bat Pest-Control Services for Cotton | USA | Plantation | *Tadarida brasiliensis mexicana* |
| 28 | Lopez-Hoffman et al. (2017) | Operationalizing the telecoupling framework for migratory species using the spatial subsidies approach to examine ecosystem services provided by Mexican free-tailed bats | US and Mexico | Plantation | *Tadarida brasiliensis* |
| 29 | Maas et al. (2013) | Bats and birds increase crop yield in tropical agroforestry landscapes | Indonesia | Plantation | Bat assemblage |

| Study nb | Authors | Article title | Country | Main habitat type | Bat species |
|---|---|---|---|---|---|
| 30 | Maine and Boyles (2015) | Bats initiate vital agroecological interactions in corn | USA | Arable land | Bat assemblage |
| 31 | Malso et al. (2017) | Chirosurveillance: The use of native bats to detect invasive agricultural pests | USA | Arable land | *Eptesicus fuscus* |
| 32 | Maslo et al. (2022) | Bats provide a critical ecosystem service by consuming a large diversity of agricultural pest insects | USA | Arable land | *Myotis lucifugus, Eptesicus fucus* |
| 33 | Manning and Ando A(2022) | Ecosystem Services and Land Rental Markets: Producer Costs of Bat Population Crashes | USA | Plantation | Bat assemblage |
| 34 | Mata et al (2021) | Combining DNA metabarcoding and ecological networks to inform conservation biocontrol by small vertebrate predators | Portugal | Mixed | Bat assemblage |
| 35 | McCracken et al. (2012) | Bats Track and Exploit Changes in Insect Pest Populations | USA | Arable land | *Tadarida brasiliensis* |
| 36 | Moiseienko and Vlaschenko (2021) | Quantitative evaluation of individual food intake by insectivorous vespertilionid bats (Chiroptera, Vespertilionidae) | Ukraine | No information | Bat assemblage |
| 37 | Montauban et al. (2021) | Bats as natural samplers: First record of the invasive pest rice water weevil Lissorhoptrus oryzophilus in the Iberian Peninsula | Spain | Arable land | *Pipistrellus pygmaeus* |
| 38 | Nguyen et al. (2019) | Vertical stratification in foraging activity of Chaerephon plicatus (Molossidae, Chiroptera) in Central Thailand | Thailand, Ban Mi District, Lopburi province | Arable land | *Chaerephon plicatus* |
| 39 | Nóbrega et al. (2023) | Insights into the habitat associations, phylogeny, and diet of Pipistrellus maderensis in Porto Santo, northeastern Macaronesia | Portugal | Mixed | *Pipistrellus maderensis* |

| Study nb | Authors | Article title | Country | Main habitat type | Bat species |
|---|---|---|---|---|---|
| 40 | Nsengimana et al. (2023) | Our good neighbors: Understanding ecosystem services provided by insectivorous bats in Rwanda | Rwanda | Arable land | Bat assemblage |
| 41 | Ocampo-Ariza et al. (2023) | Birds and bats enhance cacao yield despite suppressing arthropod mesopredation | Peru | Plantation | Bat assemblage |
| 42 | Puig-Montserrat et al. (2015) | Pest control service provided by bats in Mediterranean rice paddies: linking agroecosystems structure to ecological functions | Spain | Arable land | *Pipistrellus pygmaeus* |
| 43 | Puig-Montserrat et al. (2020) | Bats actively prey on mosquitoes and other deleterious insects in rice paddies: Potential impact on human health and agriculture | Spain | Arable land | *Pipistrellus pygmaeus* |
| 44 | Rodriguez-San Pedro et al. (2020) | Quantifying ecological and economic value of pest control services provided by bats in a vineyard landscape of central Chile | Chile | Arable land | *Myotis chiloensis, Tadarida brasiliensis* |
| 45 | Sedlock et al. (2019) | Local-Scale Bat Guild Activity Differs with Rice Growth Stage at Ground Level in the Philippines | Philippines | Arable land | Bat assemblage |
| 46 | Srilopan et al. (2018) | The wrinkle-lipped free-tailed bat (Chaerephon plicatus Buchannan, 1800) feeds mainly on brown planthoppers in rice fields of central Thailand | Thailand | Arable land | *Chaerephon plicatus* |
| 47 | Wiederholt et al. (2015) | Optimizing conservation strategies for Mexican free-tailed bats: a population viability and ecosystem services approach | USA | No information | *Tadarida brasiliensis mexicana* |
| 48 | Wiederholt et al. (2017) | Improving spatio-temporal benefit transfers for pest control by generalist predators in cotton in the southwestern US | USA | Plantation | *Tadarida brasiliensis mexicana* |

***Table S4**: List of bat species investigated in the 48 included studies and number of studies per bat species, with references supporting the bat foraging guild assigned to each species if the information was not included in the studies. For each bat species, information of foraging behaviour is provided with the corresponding reference. If foraging behaviour information was provided in the included studies, it is indicated as "Included studies".*

| Bat family | Bat species common name | Bat species | Foraging behaviour | Studies nb | Reference |
|---|---|---|---|---|---|
| Emballonuridae | Peter's Sheath-tailed Bat | *Paraemballonura atrata* | edge space aerial forager | 1 | Included studies |
| Hipposideridae | Commerson's Leaf-nosed Bat | *Hipposideros commersoni* | narrow space flutter forager | 1 | Denzinger and Schnitzler (2013) |
| Hipposideridae | Diadem Leaf-nosed Bat | *Hipposideros diadema* | narrow space flutter forager | 1 | Denzinger and Schnitzler (2013) |
| Hipposideridae | Noack's roundleaf Bat | *Hipposideros ruber* | narrow space flutter forager | 1 | Denzinger and Schnitzler (2013) |
| Hipposideridae | Pratt's Leaf-nosed Bat | *Hipposideros pratti* | narrow space flutter forager | 1 | Denzinger and Schnitzler (2013) |
| Miniopteridae | Asian Long-fingered Bat | *Miniopterus fuliginosus* | unknown | 1 | / |
| Miniopteridae | Least long-fingered Bat | *Miniopterus minor* | unknown | 1 | / |
| Miniopteridae | Major's Long-fingered Bat | *Miniopterus majori* | edge space aerial forager | 1 | Included studies |
| Miniopteridae | Manavil Long-fingered Bat | *Miniopterus manavi* | edge space aerial forager | 1 | Included studies |
| Miniopteridae | Schreibers's Long-fingered Bat | *Miniopterus schreibersii* | open space aerial forager | 2 | Included studies |
| Molossidae | Angolian Free-tailed bat | *Mops condylurus* | open space aerial forager | 1 | Included studies |
| Molossidae | Brazilian Free-tailed Bat | *Tadarida brasiliensis* | open space aerial forager | 5 | Beilke et al. (2021) |
| Molossidae | Broad-eared Free-tailed bat | *Nyctinomops laticaudatus* | open space aerial forager | 1 | Kalko et al. (2008) |
| Molossidae | European Free-tailed Bat | *Tadarida teniotis* | open space aerial forager | 1 | Included studies |
| Molossidae | Large-eared Giant Mastiff Bat | *Otomops martiensseni* | open space aerial forager | 1 | Included studies |
| Molossidae | Little Free-tailed Bat | *Chaerephon pumilus* | open space aerial forager | 1 | Bouchard (1998) Denzinger and Schnitzler (2013) |
| Molossidae | Madagascar Free-tailed Bat | *Chaerephon atsinanana* | open space aerial forager | 1 | Included studies |
| Molossidae | Malagasy White-bellied Free-tailed Bat | *Mops leucostima* | open space aerial forager | 1 | Included studies |
| Molossidae | Mexican Free-tailed bat | *Tadarida brasiliensis mexicana* | open space aerial forager | 3 | Included studies |
| Molossidae | Pallas's Mastiff Bat | *Molossus molossus* | open space aerial forager | 1 | Kalko et al. (2008) |
| Molossidae | Peter's Little Mastiff Bat | *Mormopterus jugularis* | open space aerial forager | 1 | Included studies |

| Bat family | Bat species common name | Bat species | Foraging behaviour | Studies nb | Reference |
|---|---|---|---|---|---|
| Molossidae | Southern Dog-faced bat | *Cynomops planirostris* | open space aerial forager | 1 | Included studies |
| Molossidae | Western Bonneted Bat | *Eumops perotis* | open space aerial forager | 1 | Avila-Flores et al. (2005)<br>Denzinger and Schnitzler (2013) |
| Molossidae | Wrinkle-lipped free-tailed Bat | *Chaerephon plicatus* | open space aerial forager | 1 | Sedlock et al. (2019) |
| Myzopodidae | Eastern Sucker-footed Bat | *Myzopoda aurita* | unknown | 1 | / |
| Rhinolophidae | Arcuate Horseshoe Bat | *Rhinolophus arcuatus* | narrow space flutter forager | 1 | Denzinger and Schnitzler (2013) |
| Rhinolophidae | Big-eared Horseshoe Bat | *Rhinolophus macrotis* | narrow space flutter forager | 1 | Denzinger and Schnitzler (2013) |
| Rhinolophidae | Greater Horseshoe Bat | *Rhinolophus ferrumequinum* | narrow space flutter forager | 3 | Denzinger and Schnitzler (2013) |
| Rhinolophidae | Intermediate Horseshoe Bat | *Rhinolophus affinis* | narrow space flutter forager | 1 | Denzinger and Schnitzler (2013) |
| Rhinolophidae | Least Horseshoe Bat | *Rhinolophus pusillus* | narrow space flutter forager | 1 | Denzinger and Schnitzler (2013) |
| Rhinolophidae | Lesser horseshoe Bat | *Rhinolophus hipposideros* | narrow space flutter forager | 4 | Denzinger and Schnitzler (2013) |
| Rhinolophidae | Mediterranean Horseshoe Bat | *Rhinolophus euryale* | narrow space flutter forager | 1 | Denzinger and Schnitzler (2013) |
| Vespertilionidae | African Yellow Bat | *Scotophilus dinganii* | open space aerial forager | 1 | Zachos (2020) |
| Vespertilionidae | Banana Serotine | *Afronycteris nanus* | edge space aerial forager | 1 | Zachos (2020) |
| Vespertilionidae | Bechstein's Myotis | *Myotis bechsteinii* | narrow space gleaning forager | 1 | Denzinger and Schnitzler (2013) |
| Vespertilionidae | Big Brown bat | *Eptesicus fuscus* | open space aerial forager | 2 | Beilke et al. (2021) |
| Vespertilionidae | Brown Long-eared Bat | *Plecotus auritus* | narrow space gleaning forager | 2 | Included studies |
| Vespertilionidae | Cave Myotis | *Myotis velifer* | edge space aerial forager | 1 | Li and Wilkins (2022) |
| Vespertilionidae | Chilean Myotis | *Myotis chiloensis* | edge space aerial forager | 1 | Ossa et al. (2009) |
| Vespertilionidae | Common Noctule | *Nyctalus noctula* | open space aerial forager | 1 | Denzinger and Schnitzler (2013) |
| Vespertilionidae | Common Pipistrelle | *Pipistrellus pipistrellus* | edge space aerial forager | 4 | Denzinger and Schnitzler (2013) |

| Bat family | Bat species common name | Bat species | Foraging behaviour | Studies nb | Reference |
|---|---|---|---|---|---|
| Vespertilionidae | Daubenton's Myotis | *Myotis daubentonii* | edge space trawling forager | 2 | Denzinger and Schnitzler (2013) |
| Vespertilionidae | David's Myotis | *Myotis davidii* | edge space trawling forager | 1 | Milchram et al. (2023)<br>Denzinger and Schnitzler (2013) |
| Vespertilionidae | Eastern Red Bat | *Lasiurus borealis* | edge space aerial forager | 2 | Li and Wilkins (2022) |
| Vespertilionidae | Escalera's Myotis | *Myotis escalerai* | edge space aerial forager | 1 | Zachos (2020) |
| Vespertilionidae | Eurasian Serotine | *Eptesicus serotinus* | open space aerial forager | 2 | Included studies |
| Vespertilionidae | Geoffroy's bat | *Myotis emarginatus* | narrow space gleaning forager | 1 | Zachos (2020) |
| Vespertilionidae | Gould's Wattled Bat | *Chalinolobus gouldii* | edge space aerial forager | 1 | Law and Chidel (1999) |
| Vespertilionidae | Gray long-eared Bat | *Plecotus austriacus* | narrow space gleaning forager | 2 | Included studies |
| Vespertilionidae | Hoary bat | *Lasiurus cinereus* | open space aerial forager | 1 | Included studies |
| Vespertilionidae | Japanese Pipistrelle | *Pipistrellus abramus* | edge space aerial forager | 1 | Ma et al., (2010)<br>Denzinger and Schnitzler (2013) |
| Vespertilionidae | Javan Pipistrelle | *Pipistrellus javanicus* | edge space aerial forager | 1 | Included studies |
| Vespertilionidae | Kuhl's Pipistrelle | *Pipistrellus kuhlii* | edge space aerial forager | 4 | Included studies |
| Vespertilionidae | Leisler's Noctule | *Nyctalus leisleri* | open space aerial forager | 2 | Included studies |
| Vespertilionidae | Lesser Asiatic Yellow Bat | *Scotophilus kuhlii* | edge space aerial forager | 1 | Included studies |
| Vespertilionidae | Little Brown Myotis | *Myotis lucifugus* | narrow space gleaning forager | 1 | Beilke et al. (2021) |
| Vespertilionidae | Madeira Pipistrelle | *Pipistrellus maderensis* | edge space aerial forager | 1 | Ferreira et al. (2022) |
| Vespertilionidae | Malagasy Myotis | *Myotis goudoti* | edge space aerial forager | 1 | Included studies |
| Vespertilionidae | Meridional Serotine | *Eptesicus isabellinus* | open space aerial forager | 1 | Denzinger and Schnitzler (2013),<br>Lisón et al.(2015) |
| Vespertilionidae | Nathusius's pipistrelle | *Pipistrellus nathusii* | edge space aerial forager | 1 | Included studies |
| Vespertilionidae | North American Evening Bat | *Nycticeius humeralis* | edge space aerial forager | 3 | Beilke et al. (2021) |
| Vespertilionidae | Northern Myotis | *Myotis septentrionalis* | narrow space gleaning forager | 1 | Faure et al., (1993) |
| Vespertilionidae | Orange-fingered Myotis | *Myotis rufopictus* | narrow space gleaning forager | 1 | Included studies |

| Bat family | Bat species common name | Bat species | Foraging behaviour | Studies nb | Reference |
|---|---|---|---|---|---|
| Vespertilionidae | Rufous Tube-nosed Bat | *Murina leucogaster* | unknown | 1 | / |
| Vespertilionidae | Savis's Pipistrelle | *Hypsugo savii* | edge space aerial forager | 1 | Included studies |
| Vespertilionidae | Seminole Bat | *Lasiurus seminolus* | edge space aerial forager | 1 | Beilke et al. (2021) |
| Vespertilionidae | Siver-haired bat | *Lasionycteris noctivagans* | open space aerial forager | 2 | Included studies |
| Vespertilionidae | Soprano Pipistrelle | *Pipistrellus pygmaeus* | edge space aerial forager | 4 | Included studies |
| Vespertilionidae | South-eastern Myotis | *Myotis austroriparius* | unknown | 1 | / |
| Vespertilionidae | Transparent-winged Big-eared Brown bat | *Histiotus diaphanopterus* | unknown | 1 | / |
| Vespertilionidae | Tricolored Bat | *Perimyotis subflavus* | edge space aerial forager | 2 | Li and Wilkins (2022) |
| Vespertilionidae | Western Barbastelle | *Barbastella barbastellus* | edge space aerial forager | 1 | Included studies |

**References used to classify bat foraging behaviour**

Denzinger, A. and Schnitzler, H.U. (2013). Bat guilds, a concept to classify the highly diverse foraging and echolocation behaviors of microchiropteran bats. *Frontiers in physiology* ,4, p.164.

Beilke, E.A., Blakey, R.V. and O'Keefe, J.M. (2021). Bats partition activity in space and time in a large, heterogeneous landscape. *Ecology and Evolution*, *11*(11), pp.6513-6526.

Kalko, E.K., Estrada Villegas, S., Schmidt, M., Wegmann, M. and Meyer, C.F. (2008). Flying high—assessing the use of the aerosphere by bats. *Integrative and Comparative Biology*, *48*(1), pp.60-73.

Bouchard, S. (1998). Chaerephon pumilus. *Mammalian Species*, (574), pp.1-6.

Avila-Flores, R. and Fenton, M.B. (2005). Use of spatial features by foraging insectivorous bats in a large urban landscape. *Journal of mammalogy*, *86*(6), pp.1193-1204.

Zachos, F.E. (2020). DE Wilson and RA Mittermeier (chief editors): Handbook of the Mammals of the World. Vol. 9. Bats. Lynx Edicions, Barcelona (2019). 1008 pp., 74 colour plates, 404 colour photographs, 1422 distribution maps. Hardback.€ 160, ISBN: 978–84-16,728–19-0.

Li, H. and Wilkins, K.T. (2022). Predator-prey relationship between urban bats and insects impacted by both artificial light at night and spatial clutter. *Biology*, *11*(6), p.829.

Ossa, G., Ibarra Eliessetch, J.T., Hernández, F., Gálvez Robinson, N.C., Laker, J. and Bonacic Salas, C. (2009). Preliminary acoustic analysis of Myotis chiloensis (Waterhouse, 1838), Vespertilionidae, an endemic Bat of Southern Temperate Rainforest. In *International Mammalogical Congress (10°: 2009: Mendoza, Argentina)*.

Milchram, M., Dietz, C., Mayer, F., Gurke, M., Krainer, K., Mixanig, M., Wieser, D. and Reiter, G. (2023). Moving north: Morphometric traits facilitate monitoring of the expanding steppe whiskered bat Myotis davidii in Europe. *Hystrix Ital. J. Mammal*, *34*, pp.19-23.

Law, B.S., Anderson, J. and Chidel, M. (1999). Bat communities in a fragmented forest landscape on the south-west slopes of New South Wales, Australia. *Biological Conservation*, *88*(3), pp.333-345.

Ma, J., Jones, G., Zhu, G.J. and Metzner, W. (2010). Echolocation behaviours of the Japanese pipistrelle bat Pipistrellus abramus during foraging flight. *Acta theriologica*, *55*, pp.315-332.

Ferreira, D.F., Gibb, R., López-Baucells, A., Nunes, N.J., Jones, K.E. and Rocha, R. (2022). Species-specific responses to land-use change in island insectivorous bats. *Journal for Nature Conservation*, *67*, p.126177.

Faure, P.A., Fullard, J.H. and Dawson, J.W. (1993). The gleaning attacks of the northern long-eared bat, Myotis septentrionalis, are relatively inaudible to moths. *Journal of Experimental Biology*, *178*(1), pp.173-189.

***Table S6.*** *List of species recognised as pest in Ireland, and vector of disease for Human and livestock. Lists: (1): EU priority pest; (2): Ireland protected zone plant pest; (3): Teagasc; (4): European Centre for Disease Prevention and Control. When species were categorised as pest using literature, the publication is indicated.*

| **Pest species** | **Family** | **Order** | **Type of disturbance** | **Category of pest** | **Pathogen transmitted** | **Irish distribution** | **List of pest** |
|---|---|---|---|---|---|---|---|
| *Erwinia amylovora* | Erwiniaceae | Enterobacterales | Plant Disease | Agroforestry, Forest | NONE | Present | 2 |
| *Thaumetopoea processioneata* | Noctuidae | Coleoptera | Nuisance, Plant Feeder | Forest, Human Health | NONE | Absent | 2 |
| *Tomostethus nigritus* | Tenthredinidae | Hymenoptera | Plant Feeder | Forest | NONE | Present | 2 |
| *Nysius huttoni* | Lygaeidae | Hemiptera | Plant Feeder | Agriculture | NONE | Present | 2 |
| *Bruchus rufimanus* | Chrysomelidae | Coleoptera | Plant Feeder | Agriculture | NONE | Present | 2 |
| *Cydalima perspectalis* | Crambidae | Lepidoptera | Plant Feeder | Horticulture | NONE | Present | 2 |
| *Enigmadiplosis agapanthi* | Cecidomyiidae | Diptera | Plant Feeder | Horticulture | NONE | Present | 2 |
| *Epiphyas postvittana* | Tortricidae | Lepidoptera | Plant Feeder | Agroforestry | NONE | Present | 2 |
| *Phytophthora alni* | Peronosporaceae | Peronosporales | Plant Disease | Forest | NONE | Present | 2 |
| *Phytophthora pluvialis* | Peronosporaceae | Peronosporales | Plant Disease | Forest | NONE | Present | 2 |
| *Pityogenes chalcographus* | Curculionidae | Coleoptera | Plant Feeder | Forest | NONE | Present | 2 |
| *Tuta absoluta* | Gelechiidae | Lepidoptera | Plant Feeder | Agriculture | NONE | Present | 2 |

| Pest species | Family | Order | Type of disturbance | Category of pest | Pathogen transmitted | Irish distribution | List of pest |
|---|---|---|---|---|---|---|---|
| *Sitobion avenae* | Aphididae | Hemiptera | Plant Feeder | Agriculture | Barley yellow dwarf | Present | 3 |
| *Rhopalosiphum padi* | Aphididae | Hemiptera | Plant Feeder | Agriculture | Barley yellow dwarf | Present | 3 |
| *Tipula paludosa* | Tipulidae | Diptera | Plant Feeder | Agriculture, Pasture | NONE | Present | 3, Moffat et al.,2022 |
| *Tipula oleracea* | Tipulidae | Diptera | Plant Feeder | Agriculture, Pasture | NONE | Present | 3 Moffat et al., 2022 |
| *Metopolophium dirhodum* | Aphididae | Hemiptera | Plant Feeder | Agriculture | BYDV | Present | 3 |
| *Sitona lineatus* | Curculionidae | Coleoptera | Plant Feeder | Agriculture | NONE | Present | 3 |
| *Aphis fabae* | Aphididae | Hemiptera | Plant Feeder | Agriculture | NONE | Present | 3 |
| *Psylliodes chrysocephala* | Chrysomelidae | Coleoptera | Plant Feeder | Agriculture | NONE | Present | 3 |
| *Meligethes aeneus* | Nitidulidae | Coleoptera | Plant Feeder | Agriculture | NONE | Present | 3 |
| *Acyrthosiphon pisum* | Aphididae | Hemiptera | Plant Feeder | Agriculture | NONE | Present | 3 |
| *Culicoides obsoletus* | Ceratopogonidae | Diptera | Disease Vector | Livestock | Schmallenberg virus and Blue tongue virus | Present | 4 |
| *Culicoides dewulfi* | Ceratopogonidae | Diptera | Disease Vector | Livestock | Schmallenberg virus and Blue tongue virus | Present | 4 |
| *Culicoides scoticus* | Ceratopogonidae | Diptera | Disease Vector | Livestock | Schmallenberg virus and Blue tongue virus | Present | 4 |

| Pest species | Family | Order | Type of disturbance | Category of pest | Pathogen transmitted | Irish distribution | List of pest |
|---|---|---|---|---|---|---|---|
| *Culicoides chiopterus* | Ceratopogonidae | Diptera | Disease Vector | Livestock | Schmallenberg virus and Blue tongue virus | Present | 4 |
| *Culicoides pulicaris/lupicaris* | Ceratopogonidae | Diptera | Disease Vector | Livestock | Schmallenberg virus and Blue tongue virus | Present | 4 |
| *Culicoides punctatus* | Ceratopogonidae | Diptera | Disease Vector | Livestock | Schmallenberg virus and Blue tongue virus | Present | 4 |
| *Culicoides newsteadi* | Ceratopogonidae | Diptera | Disease Vector | Livestock | Blue tongue virus | Present | 4 |
| *Culicoides impunctatus* | Ceratopogonidae | Diptera | Disease Vector | Livestock | Blue tongue virus | Present | McCarthy et al., 2016 |
| *Culicoides delta* | Ceratopogonidae | Diptera | Disease Vector | Livestock | Blue tongue virus | Present | McCarthy et al., 2016 |
| *Culicoides grisescens* | Ceratopogonidae | Diptera | Disease Vector | Livestock | Blue tongue virus | Present | McCarthy et al., 2016 |
| *Culicoides nubeculosus* | Ceratopogonidae | Diptera | Disease Vector | Livestock | Blue tongue virus | Present | McCarthy et al., 2016 |
| *Culiseta annulata* | Culicidae | Diptera | Nuisance | Livestock | Japanese encephalitis virus | Present | 4 |
| *Aedes detritus/coluzzii* | Culicidae | Diptera | Nuisance | Human Health | West Nile virus | Present | 4 |
| *Anopheles algeriensis** | Culicidae | Diptera | Disease Vector | Human Health | Malaria, | Present | Nebbak et al., 2022 Piperaki and Daikos 2016 |
| *Anopheles maculipennis complex* | Culicidae | Diptera | Disease Vector, Nuisance | Human Health | Malaria, Dirofilaria Worm, and West Nile virus | Present | 4 |

| **Pest species** | **Family** | **Order** | **Type of disturbance** | **Category of pest** | **Pathogen transmitted** | **Irish distribution** | **List of pest** |
|---|---|---|---|---|---|---|---|
| *Anopheles plumbeus* | Culicidae | Diptera | Disease Vector, Nuisance | Human Health, Livestock | Malaria | Present | 4 |
| *Coquillettidia richiardii* | Culicidae | Diptera | Disease Vector | Human Health | West Nile virus, and Dirofilarial worms | Present | 4 |
| *Culex pipiens group* | Culicidae | Diptera | Disease Vector, Nuisance | Human Health, Livestock | West Nile virus, Usutu virus, Rift Valley Fever virus, Sindbis virus, Tahyna orthobunyavirus, Dirofilarial Worms, Avian Malaria | Present | 4 |
| *Hyalomma lusitanicum* | Ixodidae | Ixodida | Ectoparasite | Livestock | NONE | Absent | 4 |
| *Ornithodorus erraticus* | Ixodidae | Ixodida | Ectoparasite | Livestock | NONE | Absent | 4 |
| *Hyalomma marginatum* | Ixodidae | Ixodida | Ectoparasite | Livestock | Crimean-Congo haemorrhagic fever virus | Absent | 4 |
| *Dermacentor reticulatus* | Ixodidae | Ixodida | Ectoparasite | Livestock | NONE | Present | 4 |
| *Rhipicephalus bursa* | Ixodidae | Ixodida | Ectoparasite | Livestock | NONE | Absent | Colebrook et al., 2004 |
| *Rhipicephalus turanicus* | Ixodidae | Ixodida | Ectoparasite | Livestock | NONE | Absent | Colebrook et al., 2004 |
| *Ixodes ricinus* | Ixodidae | Ixodida | Ectoparasite | Livestock | Borreliosis, and Tick Encephalitis | Present | 4 |

| Pest species | Family | Order | Type of disturbance | Category of pest | Pathogen transmitted | Irish distribution | List of pest |
|---|---|---|---|---|---|---|---|
| *Bovicola bovis* | Bovicoliidae | Phthiraptera | Ectoparasite | Livestock | NONE | Present | Colebrook et al., 2004 |
| *Haematopinus eurysternus* | Haematopinidae | Phthiraptera | Ectoparasite | Livestock | NONE | Present | Colebrook et al., 2004 |
| *Bovicola ovis* | Bovicoliidae | Phthiraptera | Ectoparasite | Livestock | NONE | Present | Colebrook et al., 2004 |
| *Lucilia sericata* | Calliphoridae | Diptera | Ectoparasite | Livestock | NONE | Present | Colebrook et al., 2004 |
| *Oestrus ovis* | Oestridae | Diptera | Ectoparasite | Livestock | NONE | Present | Colebrook et al., 2004 |
| *Hypoderma lineatum* | Oestridae | Diptera | Ectoparasite | Livestock | NONE | Present | Colebrook et al., 2004 |
| *Hypoderma bovis* | Oestridae | Diptera | Ectoparasite | Livestock | NONE | Present | Colebrook et al., 2004 |
| *Gasterophilus intestinalis* | Oestridae | Diptera | Ectoparasite | Livestock | NONE | Present | Colebrook et al., 2004 |
| *Ctenocephalides felis* | Pulicidae | Siphonaptera | Ectoparasite | Livestock | NONE | Present | Colebrook et al., 2004 |
| *Drosophila suzukii* | Drosophilidae | Diptera | Plant Feeder | Agriculture, Horticulture | NONE | Present | 3 |
| *Plutella xylostella* | Plutellidae | Lepidoptera | Plant Feeder | Agriculture | NONE | Present | 3 |

***Figure S2***. *Plot presenting the frequency of occurrence of pest species in the diet of each bat species across year of collection and sampling period.*

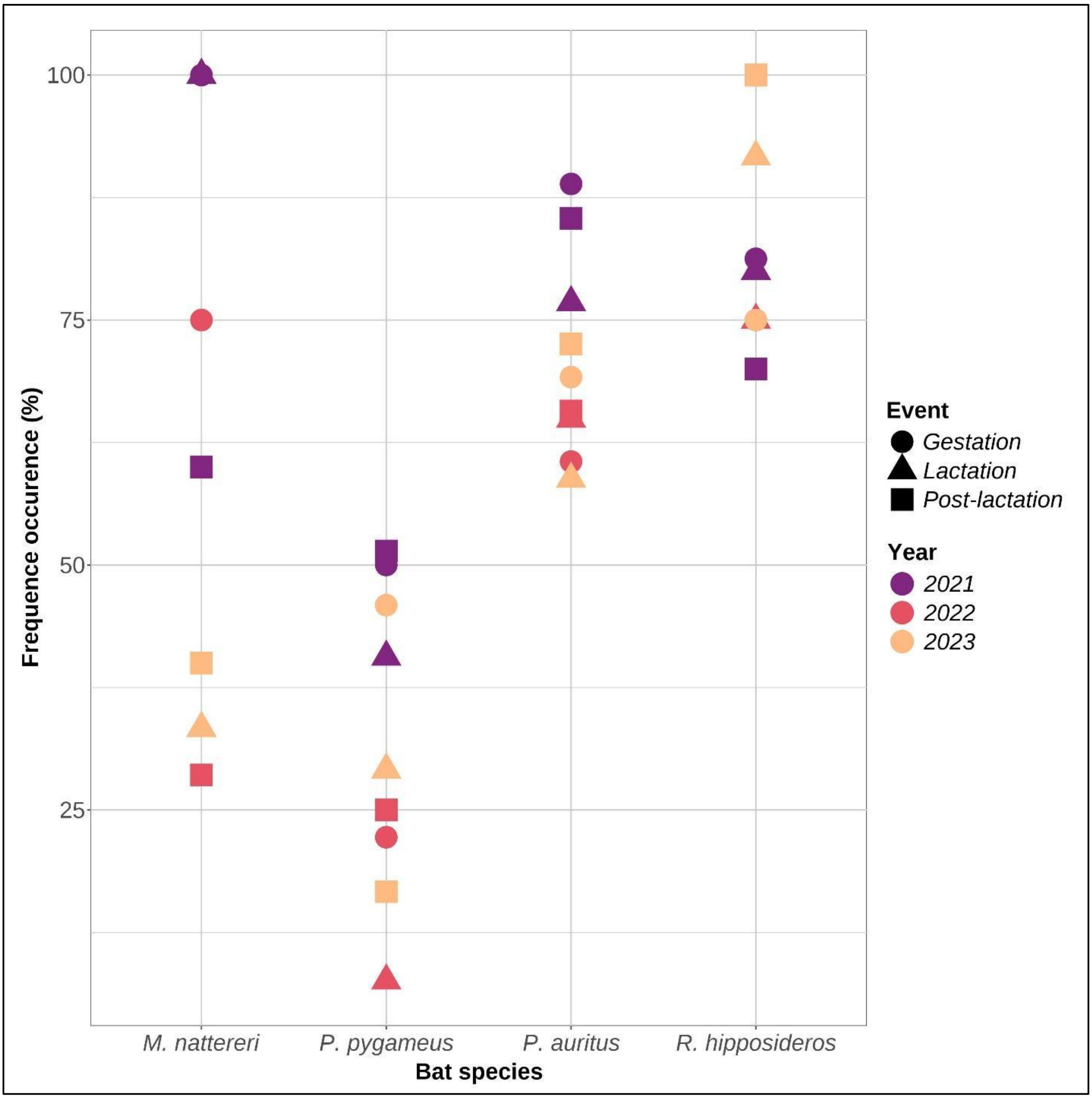

***Table S7.*** *A: Frequency of occurrence values of pest (in %) for each bat species. B: Comparison of frequencies of occurrence between bat species using Wilcoxon Mann Whitney test. Significant p-value are identified in bold.*

| ***A. Frequence of occurrence of pest across bat species*** | | | | | |
|---|---|---|---|---|---|
| ***Bat species*** | ***min*** | ***median*** | ***mean*** | ***max*** | ***SD*** |
| *Myotis nattereri* | *28.6* | *50.0* | *58.8* | *100.0* | *29.8* |
| *Pipistrellus pygmaeus* | *7.6* | *29.1* | *32.1* | *51.4* | *15.6* |
| *Plecotus auritus* | *58.8* | *69.8* | *71.4* | *88.9* | *10.5* |
| *Rhinolophus hipposideros* | *70.0* | *80.0* | *83.1* | *100.0* | *11.3* |

| ***B. Frequency of occurrence mean comparison, method: Wilcoxon Mann-Whitney pairwise*** | | | |
|---|---|---|---|
| | *Myotis nattereri* | *Pipistrellus pygmaeus* | *Plecotus auritus* |
| *Pipistrellus pygmaeus* | *0.0774* | | |
| *Plecotus auritus* | *0.3117* | ***4.10E-05*** | |
| *Rhinolophus hipposideros* | *0.0975* | ***4.40E-04*** | ***0.0417*** |

***Table S8.*** *Frequency of occurrence (FO) of each pest species identified in the diet the bat species per time point, over the three years of sampling, including median and mean.*

| | | Per time point | Over the three years | |
|---|---|---|---|---|
| species | Time point | FO (%) | FO Median (%) | FO Mean (%) |
| Agriopis aurantiaria | 2022 Gestation | 0.45 | 0.45 | 0.45 |
| Agriotes lineatus | 2023 Lactation | 0.81 | 0.81 | 0.81 |
| Agrotis ipsilon | 2021 Lactation | 0.59 | 0.88 | 1.99 |
| Agrotis ipsilon | 2022 Post-lactation | 0.44 | | |
| Agrotis ipsilon | 2023 Gestation | 0.88 | | |
| Agrotis ipsilon | 2023 Lactation | 5.65 | | |
| Agrotis ipsilon | 2023 Post-lactation | 2.4 | | |
| Agrotis segetum | 2023 Lactation | 0.4 | 0.4 | 0.4 |
| Agrotis segetum | 2023 Post-lactation | 0.4 | | |
| Anopheles algeriensis | 2022 Gestation | 0.89 | 0.89 | 0.89 |
| Arctia caja | 2022 Lactation | 0.83 | 1.29 | 1.45 |
| Arctia caja | 2023 Gestation | 1.76 | | |
| Arctia caja | 2023 Lactation | 2.82 | | |
| Arctia caja | 2023 Post-lactation | 0.4 | | |
| Argyresthia conjugella | 2022 Lactation | 0.42 | 0.41 | 0.41 |
| Argyresthia conjugella | 2023 Lactation | 0.4 | | |
| Athous haemorrhoidalis | 2022 Gestation | 0.89 | 0.66 | 0.66 |
| Athous haemorrhoidalis | 2022 Post-lactation | 0.44 | | |
| Autographa gamma | 2021 Lactation | 0.59 | 4.84 | 8.24 |
| Autographa gamma | 2021 Post-lactation | 8.42 | | |
| Autographa gamma | 2022 Gestation | 0.45 | | |
| Autographa gamma | 2022 Lactation | 1.25 | | |
| Autographa gamma | 2022 Post-lactation | 0.44 | | |
| Autographa gamma | 2023 Gestation | 11.01 | | |
| Autographa gamma | 2023 Lactation | 25 | | |
| Autographa gamma | 2023 Post-lactation | 18.8 | | |
| Celypha lacunana | 2021 Gestation | 0.59 | 0.45 | 0.61 |
| Celypha lacunana | 2021 Lactation | 1.18 | | |
| Celypha lacunana | 2022 Gestation | 0.45 | | |
| Celypha lacunana | 2022 Post-lactation | 0.44 | | |
| Celypha lacunana | 2023 Post-lactation | 0.4 | | |
| Chamaepsila rosae | 2023 Lactation | 0.4 | 0.8 | 0.8 |
| Chamaepsila rosae | 2023 Post-lactation | 1.2 | | |

| | | | | |
|---|---|---|---|---|
| Cnephasia asseclana | 2022 Lactation | 0.42 | 0.43 | 0.43 |
| Cnephasia asseclana | 2022 Post-lactation | 0.44 | | |
| Culicoides fascipennis | 2022 Post-lactation | 3.52 | 1.96 | 1.96 |
| Culicoides fascipennis | 2023 Post-lactation | 0.4 | | |
| Culicoides pulicaris | 2021 Gestation | 2.96 | 1.58 | 1.71 |
| Culicoides pulicaris | 2021 Lactation | 0.59 | | |
| Culicoides pulicaris | 2021 Post-lactation | 1.58 | | |
| Culicoides scoticus | 2021 Gestation | 0.59 | 0.88 | 0.9 |
| Culicoides scoticus | 2021 Post-lactation | 0.53 | | |
| Culicoides scoticus | 2022gestation | 0.89 | | |
| Culicoides scoticus | 2022 Post-lactation | 1.32 | | |
| Culicoides scoticus | 2023 Gestation | 0.88 | | |
| Culicoides scoticus | 2023 Lactation | 1.21 | | |
| Cydia pomonella | 2022 Post-lactation | 0.44 | 0.44 | 0.44 |
| Ditula angustiorana | 2022 Lactation | 0.42 | 0.42 | 0.42 |
| Epiphyas postvittana | 2022 Gestation | 3.57 | 1.9 | 2.25 |
| Epiphyas postvittana | 2022 Lactation | 0.83 | | |
| Epiphyas postvittana | 2022 Post-lactation | 2.2 | | |
| Epiphyas postvittana | 2023 Gestation | 0.44 | | |
| Epiphyas postvittana | 2023 Lactation | 4.84 | | |
| Epiphyas postvittana | 2023 Post-lactation | 1.6 | | |
| Hydraecia micacea | 2021 Gestation | 6.51 | 4 | 5.41 |
| Hydraecia micacea | 2021 Lactation | 7.1 | | |
| Hydraecia micacea | 2021 Post-lactation | 15.26 | | |
| Hydraecia micacea | 2022 Post-lactation | 1.32 | | |
| Hydraecia micacea | 2023 Gestation | 0.44 | | |
| Hydraecia micacea | 2023 Lactation | 3.23 | | |
| Hydraecia micacea | 2023 Post-lactation | 4 | | |
| Lacanobia oleracea | 2022 Lactation | 0.83 | 0.83 | 0.83 |
| Lyonetia clerkella | 2022 Post-lactation | 0.44 | 0.44 | 0.44 |
| Noctua pronuba | 2021 Gestation | 15.98 | 23.39 | 24.81 |
| Noctua pronuba | 2021 Lactation | 37.28 | | |
| Noctua pronuba | 2021 Post-lactation | 43.68 | | |
| Noctua pronuba | 2022 Gestation | 16.96 | | |
| Noctua pronuba | 2022 Lactation | 30.83 | | |
| Noctua pronuba | 2022 Post-lactation | 29.96 | | |
| Noctua pronuba | 2023 Gestation | 8.81 | | |

| | | | | |
|---|---|---|---|---|
| Noctua pronuba | 2023 Lactation | 23.39 | | |
| Noctua pronuba | 2023 Post-lactation | 16.4 | | |
| Orthosia cerasi | 2021 Gestation | 0.59 | 0.74 | 1 |
| Orthosia cerasi | 2022 Gestation | 0.89 | | |
| Orthosia cerasi | 2022 Lactation | 2.08 | | |
| Orthosia cerasi | 2023 Gestation | 0.44 | | |
| Otiorhynchus singularis | 2021 Gestation | 0.59 | 0.59 | 0.59 |
| Pandemis cerasana | 2021 Lactation | 0.59 | 0.44 | 0.47 |
| Pandemis cerasana | 2022 Gestation | 0.45 | | |
| Pandemis cerasana | 2023 Gestation | 0.44 | | |
| Pandemis cerasana | 2023 Lactation | 0.4 | | |
| Pandemis heparana | 2022 Lactation | 1.25 | 0.42 | 0.62 |
| Pandemis heparana | 2023 Gestation | 0.44 | | |
| Pandemis heparana | 2023 Lactation | 0.4 | | |
| Pandemis heparana | 2023 Post-lactation | 0.4 | | |
| Panolis flammea | 2021 Gestation | 0.59 | 0.49 | 0.9 |
| Panolis flammea | 2021 Lactation | 2.96 | | |
| Panolis flammea | 2021 Post-lactation | 0.53 | | |
| Panolis flammea | 2022 Gestation | 0.45 | | |
| Panolis flammea | 2022 Lactation | 0.42 | | |
| Panolis flammea | 2022 Post-lactation | 0.44 | | |
| Pasiphila rectangulata | 2022 Lactation | 0.42 | 0.42 | 0.42 |
| Peribatodes rhomboidaria | 2023 Lactation | 0.4 | 0.4 | 0.4 |
| Rhopobota naevana | 2023 Lactation | 2.42 | 2.01 | 2.01 |
| Rhopobota naevana | 2023 Post-lactation | 1.6 | | |
| Scaptomyza flava | 2022 Post-lactation | 0.44 | 0.44 | 0.44 |
| Scolytus multistriatus | 2022 Post-lactation | 0.44 | 0.44 | 0.44 |
| Sitona lineatus | 2022 Post-lactation | 0.44 | 0.44 | 0.44 |
| Tipula oleracea | 2021 Gestation | 79.88 | 35.71 | 41.15 |
| Tipula oleracea | 2021 Lactation | 55.62 | | |
| Tipula oleracea | 2021 Post-lactation | 61.58 | | |
| Tipula oleracea | 2022 Gestation | 35.71 | | |
| Tipula oleracea | 2022 Lactation | 25.83 | | |
| Tipula oleracea | 2022 Post-lactation | 26.87 | | |
| Tipula oleracea | 2023 Gestation | 51.54 | | |
| Tipula oleracea | 2023 Lactation | 10.08 | | |
| Tipula oleracea | 2023 Post-lactation | 23.2 | | |

| | | | | |
|---|---|---|---|---|
| Tipula paludosa | 2021 Gestation | 26.04 | 15.98 | 16.41 |
| Tipula paludosa | 2021 Lactation | 15.98 | | |
| Tipula paludosa | 2021 Post-lactation | 30 | | |
| Tipula paludosa | 2022 Gestation | 7.59 | | |
| Tipula paludosa | 2022 Lactation | 8.75 | | |
| Tipula paludosa | 2022 Post-lactation | 19.38 | | |
| Tipula paludosa | 2023 Gestation | 7.05 | | |
| Tipula paludosa | 2023 L:actation | 7.66 | | |
| Tipula paludosa | 2023 Post-lactation | 25.2 | | |

***Table S9.*** *Results of Wilcoxon Mann-Whitney pairwise tests comparing pest richness within bat species between year of collection and sampling period. Significant p-value (p-value < 0.05) are indicated in bold.*

| **Wilcoxon Mann-Whitney pairwise test richness across year and time collection point** | | | | | | |
|---|---|---|---|---|---|---|
| | Year | | | | Sampling period | |
| All bat species | | | | | | |
| | 2022 | 2021 | | | Gestation | Lactation |
| 2021 | **< 2e-16** | | | Lactation | 0.19574 | |
| 2023 | 0.077 | **< 2e-16** | | Post-lactation | **0.01317** | **0.00072** |
| *Plecotus auritus* | | | | | | |
| | 2022 | 2021 | | | Gestation | Lactation |
| 2021 | < 2e-16 | | | Lactation | 0.381 | |
| 2023 | **0.016** | **4.1e-12** | | Post-lactation | **7.3e-16** | **0.002** |
| *Pipistrellus pygmaeus* | | | | | | |
| | 2022 | 2021 | | | Gestation | Lactation |
| 2021 | **7.2e-6** | | | Lactation | 0.33 | |
| 2023 | **0.0206** | **0.0079** | | Post-lactation | 0.121 | 0.536 |
| *Rhinolophus hipposideros* | | | | | | |
| | 2022 | 2021 | | | Gestation | Lactation |
| 2021 | **0.0069** | | | Lactation | 0.185 | |
| 2023 | 0.4098 | **0.0238** | | Post-lactation | **0.002** | 0.105 |
| *Myotis nattereri* | | | | | | |
| | 2022 | 2021 | | | Gestation | Lactation |
| 2021 | 0.061 | | | Lactation | 0.072 | |
| 2023 | **0.016** | 0.432 | | Post-lactation | 0.149 | 1 |

***Table S10.*** *Result of* PERMANOVA tests, used to identify significant differences in pest species composition in diet between bat species. *Tables present F-statistics, $R^2$ values, degree of freedom, and p-values (Pr(>F). PERMANOVA test was performed using the 'adonis2' function of the vegan package. Significant p-values are indicated in bold. Table A: PERMANOVA test using both major and potential pest species in diet composition; Table B: PERMANOVA test using major pest species only in diet composition.*

Vegan package, adonis2 – Permutation 999

A. Major and Potential pest species

| | Df | Sum of Sqs | $R^2$ | F | Pr(>F) |
|---|---|---|---|---|---|
| Model | 3 | 34.56 | 0.0991 | 43.459 | **0.001** |
| Residual | 1185 | 314.10 | 0.9008 | | |
| Total | 1185 | 348.66 | 1.0000 | | |

B. Major Pest only

| | Df | Sum of Sqs | $R^2$ | F | Pr(>F) |
|---|---|---|---|---|---|
| Model | 3 | 12.87 | 0.8849 | 29.026 | **0.001** |
| Residual | 897 | 132.55 | 0.9115 | | |
| Total | 900 | 145.41 | 1.0000 | | |

***Table S11.*** *Frequency of occurrence in percentage (FO) of Tipula pest species in the diet of studied bat species for each maternity roosts, including minimum (min), mean, maximum (max), and standard deviation (sd).*

| **Bat species** | **Roost** | **FO min** | **FO mean** | **FO max** | **FO sd** |
|---|---|---|---|---|---|
| *Myotis nattereri* | Ennisnag | 0.00 | 24.00 | 71.43 | 29.32 |
| | Glengarriff | 40.00 | 72.50 | 100.00 | 32.02 |
| | Letterfrack | 33.33 | 33.33 | 33.33 | NA |
| *Pipistrellus pygmaeus* | Dromore wood | 5.00 | 15.83 | 60.00 | 21.78 |
| | Glendalough | 0.00 | 3.06 | 12.50 | 4.97 |
| | Kilafin | 0.00 | 0.00 | 0.00 | 0.00 |
| | Birr | 7.69 | 52.09 | 100.00 | 37.72 |
| *Plecotus auritus* | Kinvarra | 25.00 | 82.22 | 100.00 | 25.14 |
| | Carra James | 35.00 | 81.67 | 100.00 | 21.65 |
| | Ennisnag | 0.00 | 19.14 | 60.00 | 23.28 |
| | Glengarriff | 0.00 | 27.24 | 88.89 | 29.45 |
| | Inagh | 0.00 | 46.06 | 100.00 | 37.05 |
| | Kilmore | 6.25 | 57.04 | 100.00 | 36.19 |
| | Miltown | 15.00 | 52.28 | 100.00 | 33.21 |
| | Letterfrack | 0.00 | 22.81 | 65.00 | 24.36 |
| *Rhinolophus hipposideros* | Kinvarra | 66.67 | 79.21 | 100.00 | 11.45 |
